# Gut-specific H3R signaling orchestrates microglia-dependent resolution of peripheral inflammation

**DOI:** 10.1101/2024.07.11.603031

**Authors:** Kerstin Dürholz, Mathias Linnerbauer, Eva Schmid, Heike Danzer, Lena Lößlein, Lena Amend, Leona Ehnes, Michael Frech, Vugar Azizov, Fabian Schälter, Arne Gessner, Sébastien Lucas, Till-Robin Lesker, R. Verena Taudte, Jörg Hofmann, Felix Beyer, Hadar Bootz-Maoz, Yasmin Reich, Hadar Romano, Daniele Mauro, Ruth Beckervordersandforth, Wei Xiang, Aiden Haghikia, Cezmi A. Akdis, Francesco Ciccia, Tobias Bäuerle, Kerstin Sarter, Till Strowig, Nissan Yissachar, Georg Schett, Veit Rothhammer, Mario M. Zaiss

**Affiliations:** Department of Internal Medicine 3, Rheumatology and Immunology, Friedrich-Alexander-Universität Erlangen-Nürnberg (FAU) and Universitätsklinikum Erlangen, Erlangen, Germany; Deutsches Zentrum Immuntherapie (DZI), Friedrich-Alexander-Universität Erlangen-Nürnberg (FAU) and Universitätsklinikum Erlangen, Erlangen, Germany; Department of Neurology, Universitätsklinikum Erlangen, Friedrich-Alexander-Universität Erlangen-Nürnberg (FAU), 91054 Erlangen, Germany; Department of Microbial Immune Regulation, Helmholtz Centre for Infection Research, Braunschweig, Germany; Hannover Medical School, Hannover, Germany; Institute of Experimental and Clinical Pharmacology and Toxicology, Friedrich-Alexander University (FAU) Erlangen-Nürnberg, Erlangen, Germany; Core Facility for Metabolomics, Philipp University Marburg, Marburg, Germany; Department of Biology, Division of Biochemistry, Friedrich-Alexander University (FAU), 91058 Erlangen, Germany; Institute of Biochemistry, Friedrich-Alexander University of Erlangen-Nürnberg, Erlangen, Germany; The Goodman Faculty of Life Sciences, and Bar-Ilan Institute of Nanotechnology and Advanced Materials, Bar-Ilan University, Ramat-Gan, 5290002, Israel; Department of Precision Medicine, University of Campania "Luigi Vanvitelli", Naples, Italy; Department of Molecular Neurology, University Hospital Erlangen, Friedrich-Alexander-Universität Erlangen-Nürnberg (FAU), Germany; Department of Molecular Cell Biology, Institute of Biochemistry and Pathobiochemistry, Ruhr University Bochum, Bochum 44801, Germany; Swiss Institute of Allergy and Asthma Research (SIAF), University of Zurich, Davos, Switzerland; Radiologisches Institut, Universitätsklinikum Erlangen, Friedrich-Alexander-Universität Erlangen-Nürnberg, Erlangen, Germany

**Keywords:** histamine, microglia, gut-joint axis, gut-brain axis, gut-CNS-joint axis, resolution

## Abstract

Chronic inflammatory diseases, like rheumatoid arthritis (RA) have been described to cause central nervous system (CNS) activation. Less is known about environmental factors that enable the CNS to suppress peripheral inflammation in RA. Here, we identified gut microbiota-derived histamine as such factor. We show that low levels of histamine activate the enteric nervous system, increase inhibitory neurotransmitter concentrations in the spinal cord and restore homeostatic microglia, thereby reducing inflammation in the joints. Selective histamine 3 receptor (H3R) signaling in the intestine is critical for this effect, as systemic and intrathecal application did not show effects. Microglia depletion or pharmacological silencing of local nerve fibers impaired oral H3R agonist-induced pro-resolving effects on arthritis. Moreover, therapeutic supplementation of the SCFA propionate identified one way to expand local intestinal histamine concentrations in mice and humans. Thus, we define a gut-CNS-joint axis pathway where microbiota-derived histamine initiates the resolution of arthritis via the CNS.

**Graphical Abstract:** 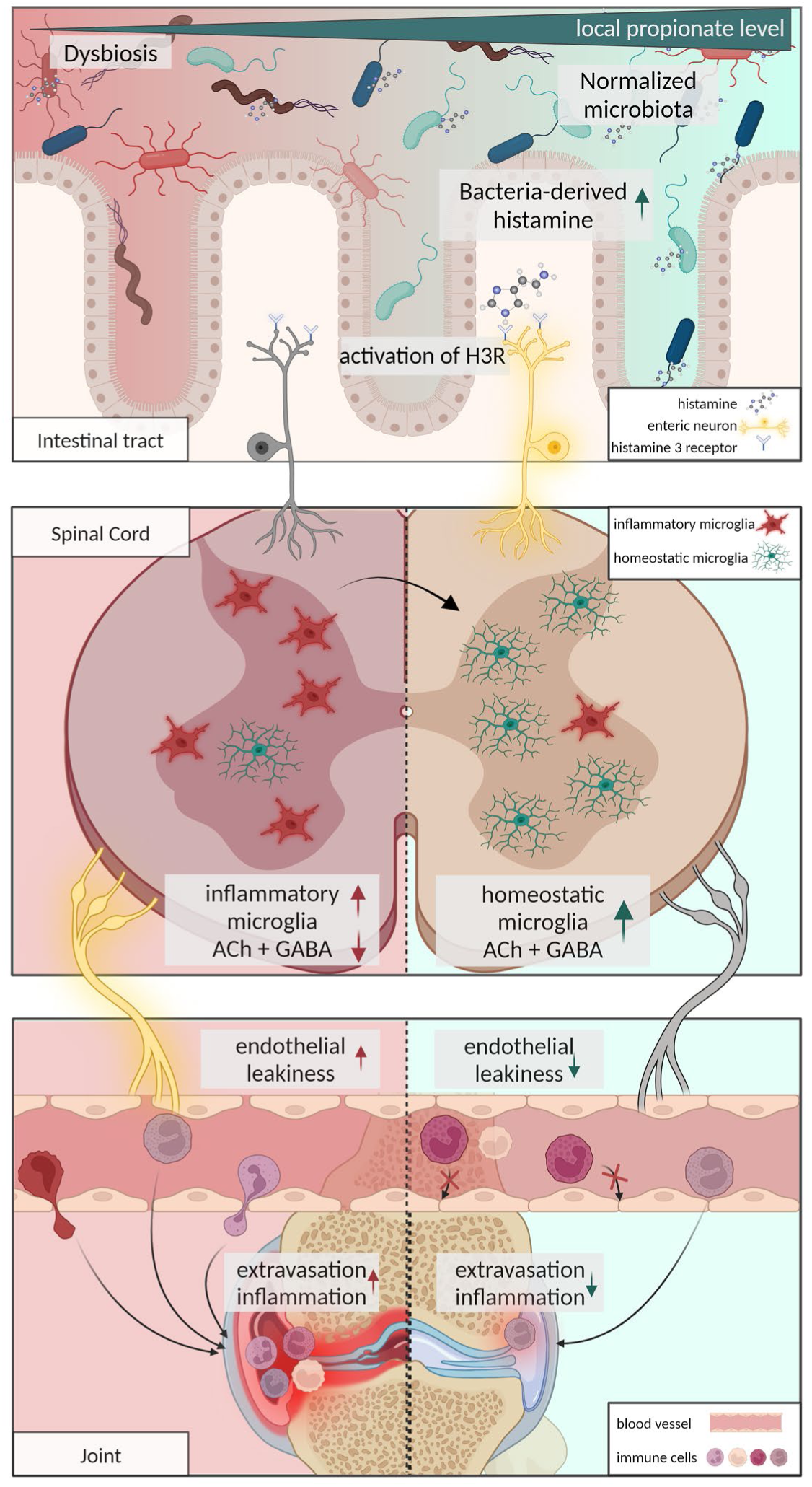

- Gut microbiota-derived histamine activates enteric neurons via H3R
- Local intestinal H3R activation induces shift to homeostatic microglia in the spinal cord
- CNS controlled decrease in endothelial leakiness resolves synovial inflammation

## Introduction

Rheumatoid arthritis (RA) is one of the most common and severe chronic inflammatory diseases with a lack of spontaneous resolution, which generally requires lifelong treatment with anti-rheumatic drugs. The disease is characterized by chronic synovial inflammation that leads to the destruction of the affected joints and increases disability (1). One essential step to initiate the resolution of arthritis is to prevent the influx of new cells via trans-endothelial migration from blood vessels into the inflamed synovium while at the same time draining pro-inflammatory cells leave the affected joints. Earlier studies showed the potential impact of the central nervous system (CNS) and sympathetic nerve fibers in control of blood flow and vascular permeability in the arthritic joints (2, 3) and its direct relevance for resolution (4, 5). However, little effort has been made to unravel how these processes are controlled and whether and how environmental factors, such as the microbiota, can initiate resolution of arthritis via the CNS. That is surprising given the vast literature on microbial dysbiosis (6, 7), the mucosal origin hypothesis (8), or the gut-joint axis (9) and its significant role in RA.

Beyond joint pathology, RA is associated with neuropsychiatric comorbidity including depression (10), anxiety (11), and an increased risk of developing neurodegenerative diseases in later life (12). Pre-clinical studies from the early 90s (13, 14), and 2000s (15-17) highlighted the potential neuro-immune crosstalk in animal models for RA by showing that peripheral synovial inflammation is closely linked to the CNS.

In the CNS, the biogenic vasoactive amine histamine, which is known for its acute immediate hypersensitivity responses, also acts as a neurotransmitter in the histaminergic system. Histamine was shown to have pleiotropic effects on immune cells that are dependent on signalling via one of its four receptors (H1R-H4R). In preclinical models of arthritis, cellular-derived histamine was so far described to exert pro-arthritic effects (18) mainly via H4R expression on hematopoietic cells (19, 20) and by stimulating osteoclastogenesis (21). In contrast, H3R is expressed by cells of the CNS and histaminergic neurons but also on intestinal endocrine cells (22) such as enterochromaffin cells (23) linking the gut to the CNS (24). Recent studies using new human H3R ligands revealed anti-inflammatory effects of H3R activation on neuroinflammation (25), matching reports of exacerbated of neuroinflammatory disease in H3R knockout mice (26).

In addition to mammalian cells, bacteria can also secrete histamine following decarboxylation of histidine via the enzyme histidine decarboxylase (HDC) (27). However, the influence of microbiota-derived histamine on host immunological processes, i.e. the resolution of inflammation is yet poorly understood (28).

Here, we exploited untargeted metabolomics and mRNA sequencing of microglia from spinal cord tissues after oral histamine treatment in collagen-induced arthritis (CIA) or experimental autoimmune encephalomyelitis (EAE) mice to identify potential pro-resolving mediators of synovial and central inflammation; assessed their function *in vivo* using PLX-5622-driven microglia or QX-314 + bupivacaine-induced nerve fiber blockage; and extended these findings to functional magnetic resonance tomography (MRT) analysis for vascular permeability in synovial tissues. The resulting data emphasizes the importance of an acutely induced inflammatory microglial phenotype for arthritis persistence, which can be restored to a homeostatic microglial gene expression profile by microbiota-derived histamine, resulting in rapid resolution of synovial inflammation.

Taken together, these data significantly contribute to our understanding of the gut-joint axis by implicating CNS microglial cells as a direct mediator of inter-organ signaling, thereby providing new therapeutic options to resolve arthritis. Further, fiber-rich or short-chain fatty acid (SCFA) propionate treatments naturally increased microbiota-derived histamine levels in mice as well as RA and multiple sclerosis (MS) patients. These are important findings as they imply the value of minimally invasive treatments, such as diet and supplementation in resolution of inflammation.

## Results

### Propionate-induced microbial metabolites transfer pro-resolving effects

Prophylactic nutritional fiber or propionate supplementation was shown effective in RA and MS animal models (29, 30). To study potential therapeutic effects, we supplemented the drinking water of CIA mice from the peak of disease at 30 dpi with 150mM propionate (C3) or high fiber supplementation. Twenty days following C3 or high fiber supplementation CIA mice showed significantly improved clinical signs of arthritis (**Fig. 1a** and **Suppl. Figure 1a)**, reduced splenic Th17 cells (**Fig. 1b**), and restored systemic bone density compared to respective non-treated controls (**Fig. 1c and Suppl Fig. 1a**). C3 treatment increased total SCFA concentrations in the intestine (**Fig. 1d**) and reestablished a gut microbial composition similar to the pre-clinical phase in healthy controls by reducing previously identified arthritis-related species such as from the *Akkermansiaceae (31)* and *Enterobacteriaceae (32)* family while increasing the Shannon index over non-C3-treated-CIA control mice. (**Fig. 1e**). To address whether C3-modulated microbiota contributes to this finding, we established a fecal microbiota (FM) transplantation model in CIA mice (**Fig. 1f**). To that end, FM donor DBA/1 mice were supplemented with C3 in the drinking water for 3 weeks. FM (C3 FM) was harvested and further divided into FM pellet (C3 pellet) and FM supernatant (C3 supernatant) following centrifugation (**Fig. 1g**). Similar to therapeutic C3 supplementation in the drinking water, C3 FM transfer, when done at the peak of arthritis, promoted fast resolution compared to transplantation of control FM (**Fig. 1h**). In contrast to C3 supplementation alone, C3 FM was superior in accelerating the resolution of synovial inflammation from 20 to 5 days after the start of the respective intervention (**Fig. 1h**). Within the three FM transfer groups, only C3 supernatant was as effective as complete C3 FM, while the C3 pellet fraction showed no statistical clinical improvement (**Fig. 1h**). This finding was reflected in the flow cytometry analyses of spleen cells, which showed that splenic Th17 cells decreased fast after C3 FM treatment as well as after supplementation of C3 in the drinking water (**Supplementary Figure 1b**). Comparison of 16S rRNA bacterial community profiles following C3 FM transfer revealed clear differences in the β-diversity 5 days after transfer over FM controls (**Fig. 1i**) and an enrichment in *Lactobacillaceae* accompanied by lower levels of *Lachnospiraceae* (**Fig. 1j, k**). The effectiveness of *Lactobacillaceae* as probiotics, especially for *Lactobacillus johnsonii,* which was increased in our experiments, has been shown in a variety of inflammatory diseases (33-35).

**Figure 1:**
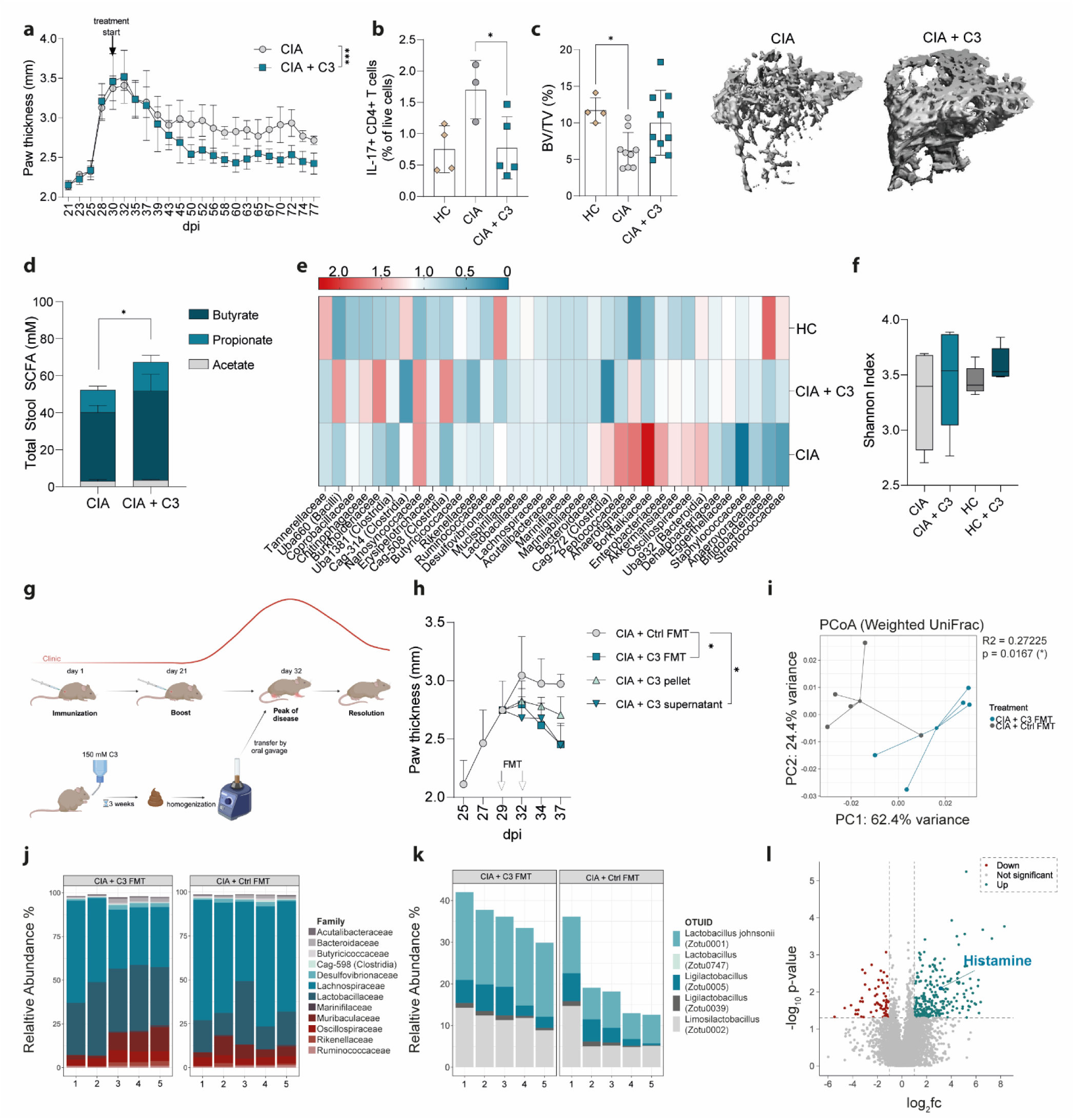
Propionate-induced microbial metabolites transfer pro-resolving effects. **a.** Clinical arthritis score shown as paw thickness (mm) of CIA mice ± 150 mM C3 (n=4-5) in the drinking water starting 30 dpi **b.** Flow cytometric analysis of IL-17+ CD4+ T cells in the spleen of healthy mice and CIA mice ± C3**c.** quantification of bone mass (BV/TV) and representative µCT images of the trabecular part of tibial bone of healthy mice and CIA mice ± C3, analyzed 77 dpi **d.** SCFA levels in the cecum content of CIA mice ± C3 **e.** 16s rRNA sequencing of the cecum content of healthy mice, CIA mice and CIA mice ± C3 in the drinking water **f.** Alpha-diversity measure of 16S rRNAseq data **g.** Experimental layout of FMT experiment. This overview was generated with BioRender **h.** Clinical arthritis score shown as paw thickness (mm) of CIA mice treated with FMT of naïve donors, C3-treated donors, supernatant of C3-treated donors or pellet of C3-treated donors **i.** PCoA Plot of 16S rRNA sequencing of mice after control or C3 FMT **j.** Relative abundance of the bacterial families identified by 16S rRNA sequencing. Permutational multivariate analysis of variance (ADONIS) was significant (R2=0.27225, P = 0.0167(*)) **k.** Relative abundance of most strongly changed Lactobacillaceae strains **l.** Volcano Plot of untargeted metabolomics analysis of stool supernatant fraction obtained from control vs C3-treated FMT donors, Cut-off were p-value < 0.05 and -1 < log2fc < 1. Data are expressed as the mean ± sd. Statistical difference was determined by ADONIS (l), One-way ANOVA (b,c,f),Student’s t-test (d) and Area under the curve (a, h). *p < 0.05; **p < 0.01; ***p < 0.001; ****p < 0.0001. CIA = collagen-induced arthritis, C3 = Propionate, dpi = days post immunization, FMT = fecal microbiota transfer.

Because C3 supernatant FM transfer was equally effective as complete C3 FM, we next performed untargeted metabolic analyses to identify possible effector molecules induced by C3 nutritional supplementation. Volcano plots of C3 supernatant FM identified significant up-, and downregulated metabolites over FM control supernatant (**Fig. 1l**). Next, to identify responsible effector molecules in C3 supernatant FM, size exclusion chromatography (SEC) was used to separate the supernatant into 5 sub-fractions with different ranges of molecular size. Strikingly, only C3 supernatant FM fraction number 1, containing only the smallest molecules, showed similar pro-resolving effects as unfractionated C3 supernatant FM (**Supplementary Figure 1c and 1d**). Untargeted metabolomics analysis of the non-fractioned C3 supernatant identified histamine among the most upregulated metabolites, which is also part of the pro-resolving Fraction 1, due to its small molecular weight of 111.15 Dalton (**Fig. 1k**). Another report also showed a tendency for histamine levels to increase in the colon after C3 treatment (36). Together, all these observations suggest potent peripheral pro-resolving effector functions of intestinal histamine that is locally increased in the intestine following high fiber or C3 dietary supplementation in mice.

### Resolution of arthritis depends on intestinal histamine and H3R signaling

We then attempted to identify the metabolite responsible for that prompt pro-resolving effect to continue with targeted mechanistic analyses. Therefore, CIA mice (30 dpi) were orally treated with histamine at slightly higher concentrations as identified in histamine ELISA of stool extracts from the MS and RA patient samples (**Supplementary Figure 2a)**. Oral histamine treatment at the peak of disease rapidly improved clinical arthritis scores in CIA mice within 5 days of treatment (**Fig. 2a**). Histological analysis of hind paws confirmed this observation showing reduced inflammatory lesions in the joints of histamine-treated CIA mice (**Fig. 2b**). Multiplex cytokine and chemokine immunoassays of respective sera revealed specifically differences in CCL5 and CCL2 immune cell chemotactic cytokines, whereas other mediators remained unchanged (**Fig. 2c** and **Supplementary Figure 2b-t**). Further, β-diversity analysis together with taxonomic profiling of the bacterial community following 16S rRNA amplicon sequencing identified changed intestinal microbiota compositions following oral histamine treatment, again with an increased relative abundance of *Lactobacillaceae* (37) (**Fig. 2d-f**). Furthermore, members of this family, such as *Lactobacillus reuterii* have been described as histamine producers in the intestine. Histamine produced by *Lactobacillus* inhibited proinflammatory cytokine production (38). This led us to investigate if microbial-secreted histamine would have similar pro-resolving effects as orally supplemented histamine on synovial inflammation. Recently, Barcik et al. (39) developed an *Escherichia coli* BL21 (*E.coli*) strain that was genetically modified to express the *Morganella morganii* derived histidine decarboxylase enzyme (HDC) (*E.coli* HDC^+^), which is responsible for catalyzing the decarboxylation of histidine to histamine. Oral transfer of *E.coli* HDC^+^ at the peak of disease significantly reduced arthritis in CIA mice compared to *E.coli*-treated or untreated CIA control mice (**Fig. 2g**).

**Figure 2:**
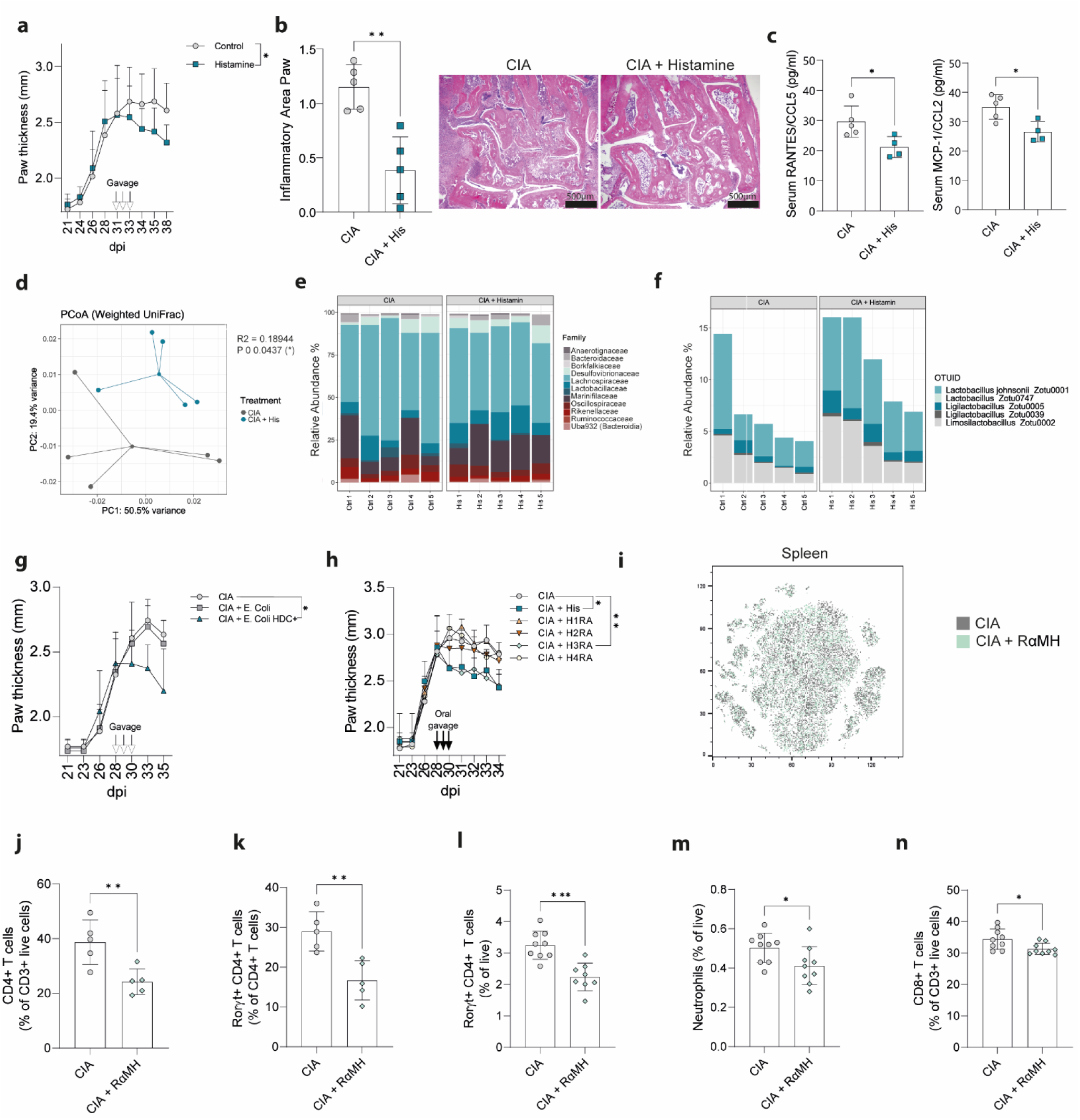
Resolution depends on intestinal histamine and H3R signaling. **a.** Clinical arthritis score shown as paw thickness (mm) of CIA mice ± histamine (n=12-14) at peak of disease **b.** Inflammatory Area quantification from H&E stained paw sections of CIA mice ± histamine (n = 5) and exemplary histology pictures **c.** Serum levels of RANTES/CCL5 and MCP-1/CCL2 **d.** PCoA Plot of 16S rRNA sequencing of CIA mice ± histamine. Permutational multivariate analysis of variance (ADONIS) was significant (R2=0.18944, P =0.0437(*)) **e.** Relative abundance of the bacterial families identified by 16S rRNA sequencing **f.** Relative abundance of most changed Lactobacillaceae strains after histamine treatment **g.** Clinical arthritis score shown as paw thickness (mm) of CIA mice after transfer of PBS (Control), E. Coli or HDC+ E. Coli at peak of disease (n = 4-5) **h.** Clinical arthritis score shown as paw thickness (mm) of CIA mice after oral transfer of Histamine or specific agonists for H1R-H4R. (n = 4-5) **i.** t-SNE Plot of spectral flow cytometric analysis of the spleen from CIA mice + H3R agonist RαMH **j.** CD4+ cells in the synovium **k.** RORγt+ CD4+ T cells in the synovium **l.** RORγt+ CD4+ T cells in the pLN **m.** Neutrophils in the pLN **n.** CD8+ T cells in the pLN. Data are expressed as the mean ± sd. Statistical difference was determined by Student’s t-test (b, c, j-n), t-test or one-way ANOVA of area under the curve (a, g, h), and ADONIS (d). *p < 0.05; **p < 0.01; ***p < 0.001; ****p < 0.0001. dpi = days post immunization, CIA = collagen-induced arthritis, pLN = popliteal lymph node.

Histamine was shown to exert its effects via four different histamine receptors H1R-H4R. These receptors differ in their affinity for histamine, with H1R and H2R having a lower affinity and H3R and H4R having a higher affinity (40). Oral treatment of CIA mice with selective H1R-H4R agonists at peak of disease (30 dpi) identified that only the H3R agonist replicated the strong pro-resolving effects of histamine itself (**Fig. 2h**). This pro-resolving effect was independent from the type of H3R agonist used, as both R(-)-alpha-methylhistamine dihydrochloride (RαMH) (41) and Immethridine dihydrobromid (IDHB) (42) showed similar pro-resolving effects (**Supplementary Figure 3a**). To confirm that only a site-specific increase of histamine in the intestine is initiating the resolution of arthritis, we compared intraperitoneal (i.p.) and intrathecal (i.th.) to oral treatment with H3R agonist in CIA mice (**Supplementary Figure 3a, b**). Only oral delivery showed significant pro-resolving effects on arthritis along with increased intestinal length (**Supplementary Figure 3c**), as an indicator for reduced intestinal inflammation. Moreover, with increasing H3R agonist concentrations, pro-resolving effects disappeared and even exacerbated clinical arthritis scores were observed (**Supplementary Figure 3d**). While t-distributed stochastic neighbor embedding (t-SNE) plots of flow cytometry analysis in spleen did not show differences in cell clustering with H3R agonist treatment (**Fig. 2i)**, CD4^+^ T cells and Th17 cells decreased in the synovial tissue (**Fig. 2j, k)** and Th17 cells, neutrophils and CD8^+^ T cells decreased in the draining popliteal lymph node (pLN) after H3R agonist treatment (**Fig. 2l-n**). As previously shown by others and us, a key mechanistic contributor to the anti-inflammatory effect of C3 is the induction of regulatory T cells (Treg) (43, 44). However, H3R agonist treatment does not induce regulatory T cells (Supplementary Figure 3e). Therefore, the pro-resolving mechanism of C3-induced intestinal histamine is independent of the classical C3-associated Treg induction. Together, these results demonstrated low-level histamine concentrations in the intestine promote the resolution of arthritis via local H3R signaling.

### H3R agonist stimulates the enteric neural system in arthritic mice causing anti-inflammatory responses

Our results show, that low dosage bacteria-derived histamine induces resolution of inflammation via specific H3R activation in the intestinal tract. H3R is mostly expressed on cells of the nervous system. In the intestine specifically, expression is interestingly limited to cells of the enteric nervous system (ENS) (45) and a very low number of endocrine cells (22). To assess H3R induced changes in the ENS, we applied an *ex vivo* gut organ culture system (46) that maintains tissue architecture, yet allows tight experimental control to perform whole-mount staining on the myenteric plexus after H3R agonist stimulation (**Fig. 3a**). The *c-fos* gene is induced by a broad range of stimuli, and has been commonly used as a reliable marker for neuronal activity (47). Histological mean fluorescence intensity (MFI) quantification in wildtype *ex vivo* gut organ cultures following H3R agonist (RαMH) treatment revealed increased *cFOS* nuclear localization in βIII-Tubulin (*Tuj1*)-positive myenteric neurons (**Fig. 3b**), consistent with previously published results by Breunig et al showing increased excitation of enteric neurons through H3R activation (48).

**Figure 3:**
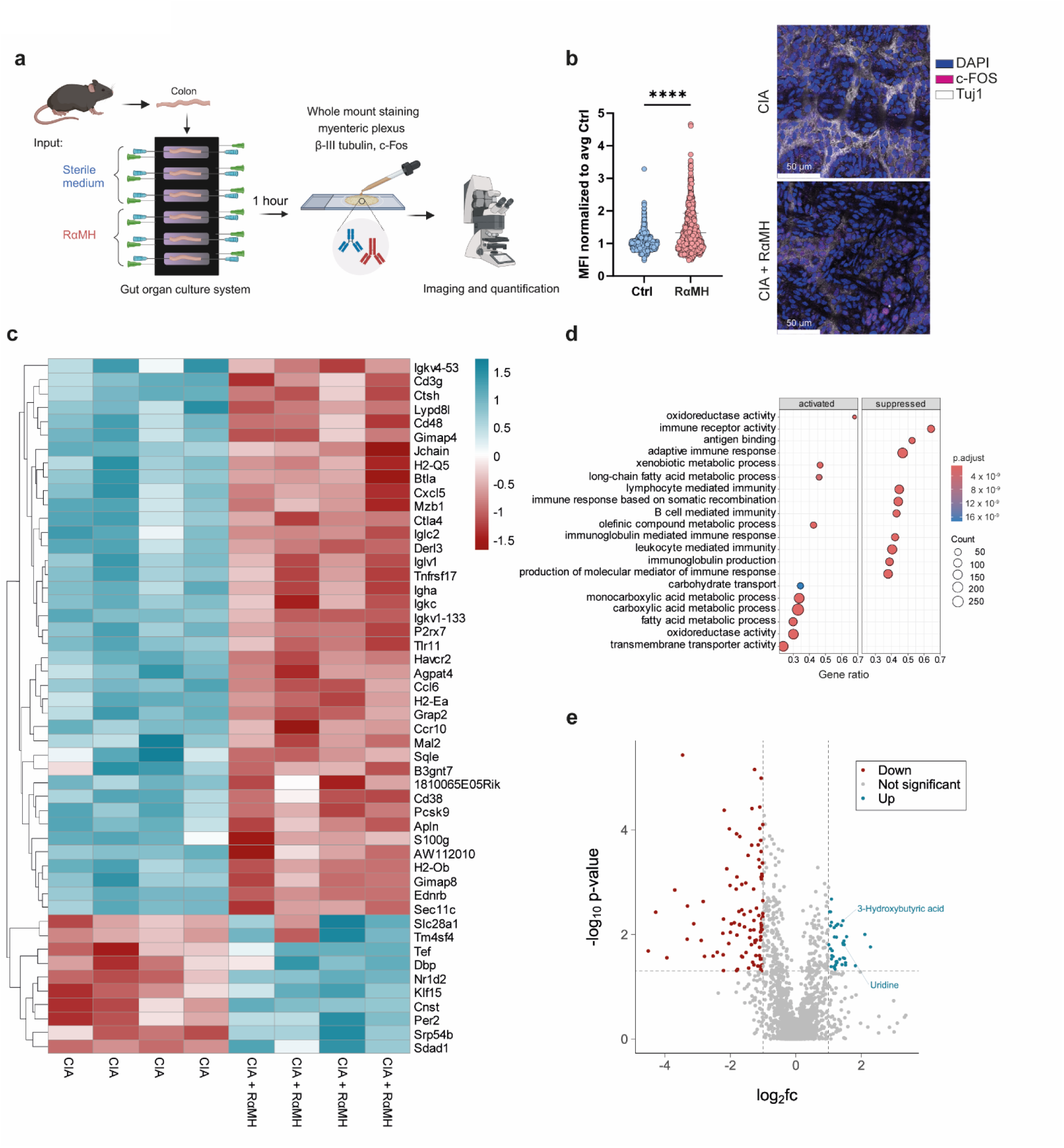
H3R agonist stimulates the ENS in arthritic mice causing anti-inflammatory responses. **a.** Experimental layout of *ex vivo* intestinal organ culture system. This overview was generated with BioRender **b.** normalized MFI of c-FOS in βIII-Tubulin+ enteric neurons after RαMH stimulation and exemplary pictures **c.** Heatmap of the top 50 regulated genes identified by RNAseq of intestinal tissue of CIA mice after in vivo RαMH treatment **d.** Gene set enrichment analysis (GSEA) of intestinal RNAseq data **e.** Volcano plot of untargeted metabolomics data of serum of CIA mice ± RαMH. Cut-off were p-value < 0.05 and -1 < log2fc < 1. Data are expressed as the mean ± sd. Statistical difference was determined by Student’s t-test. ****p < 0.0001.

Bulk RNAseq experimental data from intestinal tissues of *in vivo* H3R agonist (RαMH) treated CIA mice revealed significant differences in gene expression profiles over non-treated CIA mice (**Fig. 3c**). Gene ontology analysis showed that oral H3R agonist treatment of CIA mice induced a prominent inflammation suppressing phenotype in the intestinal tissue (**Fig. 3d**). The most suppressed genes were associated with adaptive immune response (CD3g, Jchain, Btla, Tnfrsf17), lymphocyte mediated immunity (Iglc2) and B cell mediated immunity (IL7R, Igha) (**Supplementary Figure 4)**. These findings were further supported by untargeted metabolomics analysis of serum metabolites (**Fig. 3e**). Metabolites, which were increased upon H3R stimulation, were uridine and 3-Hydroxybutyric acid, which are not only known for their anti-inflammatory properties (49, 50) but also for their role in neuroprotection (51, 52) and suppression of microglia responses (53). Taken together these data suggest, that local H3R activation in the intestine induces an anti-inflammatory milieu, and increases the neuronal activity of enteric neurons.

### Microglia depletion impairs pro-resolving effects of histamine

To formally address the contribution of the central nervous system in histamine-mediated resolution of arthritis, we next characterized cellular changes in the CNS followed by respective *in vivo* cell depletion assays. Peripheral inflammation increases *p38* phosphorylation in neurons and microglia, especially in the dorsal horn in the spinal cord (54, 55). As demonstrated by Boyle et al., local inhibition of p38 phosphorylation in the spinal cord did suppress arthritis(15). Analysis of the lumbar area (L3-L6) of the spinal cord (**Fig. 4a**) that innervates the paws (56), following oral H3R agonist treatment in CIA mice showed reduced *c-fos* and *p38* phosphorylation (**Fig. 4b, c**). Spectral flow cytometry multiparameter analysis of spinal cord single cells revealed significant cellular changes following H3R agonist treatment in CIA mice (**Fig. 4d**), most prominent within the CD86^+^ microglial cell fraction that was restored to the levels observed in healthy control (HC) mice (**Fig. 4e**). Further microglial characterization revealed a shift from inflammatory back to TMEM119^+^ homeostatic microglia (**Fig. 4f**). Interestingly, when we treated microglia, astrocytes or neurons directly with H3R agonist *in vitro*, we could not observe lower levels of inflammatory gene expression or of CCL2 and CCL5 chemokine secretion (**Supplementary Figure 5)**. When applying oral H3R agonist treatment around peak disease in the EAE animal model for MS, where microglial cells were shown to promote inflammation (57, 58), we significantly attenuated clinical scores and reduced inflammatory microglial cells in the spinal cord (**Supplementary Figure 6**) as seen in CIA mice. Next, we investigated the influence of microglia in CIA mice using the colony stimulating factor 1 receptor inhibitor, PLX5622 (PLX), to deplete the microglia population (58-60). Following short-term PLX treatment (25–30 dpi) in CIA mice, the prominent H3R agonist-mediated pro-resolving effects were lost when microglia were depleted (**Fig. 4g**). Taken together, these results show that H3R activation in the intestine results in a phenotypic switch from pro-inflammatory to homeostatic microglial cells in the spinal cord, which are essential for the H3R induced resolution of arthritis.

**Figure 4:**
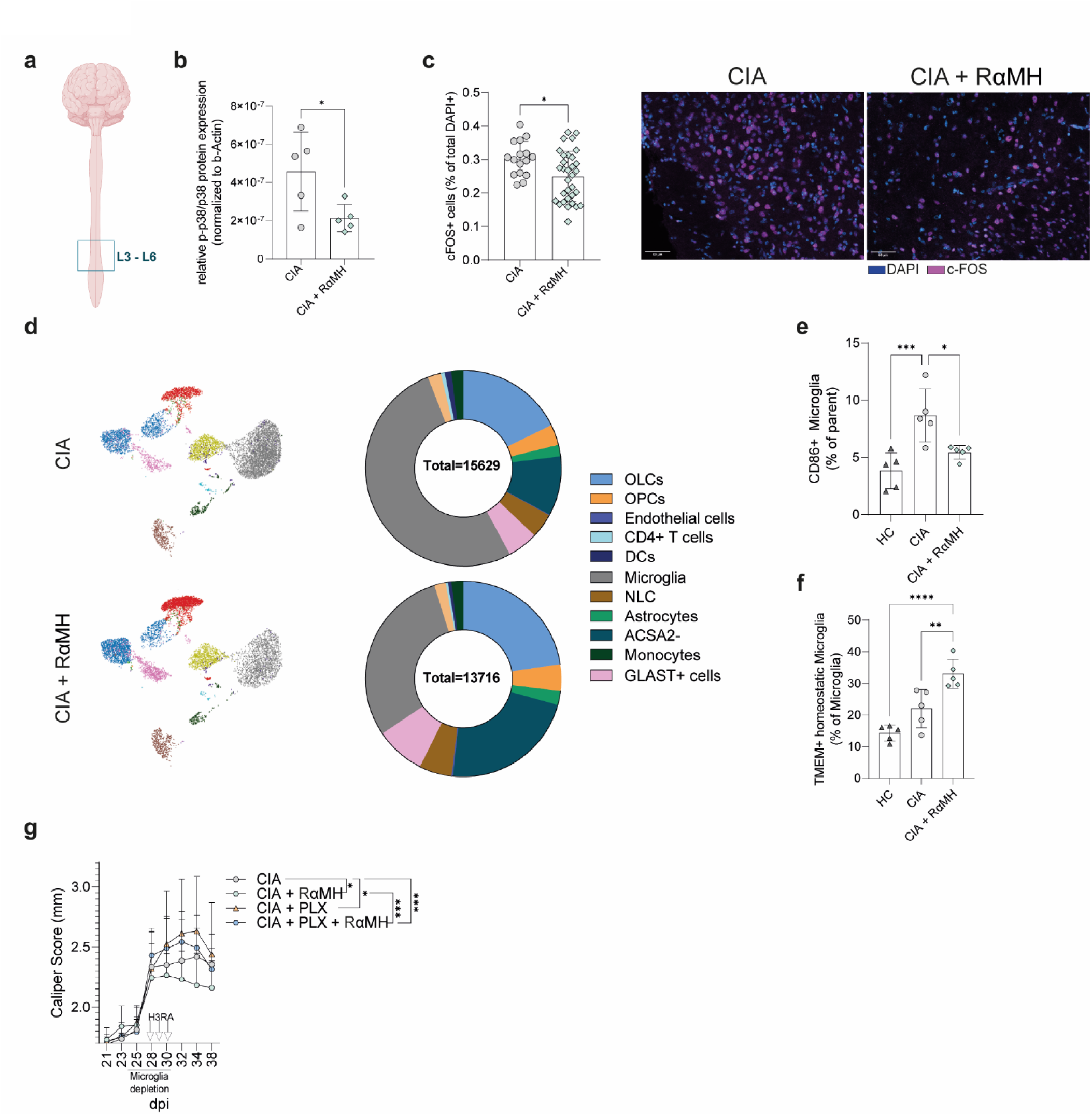
Microglia depletion impairs histamine’s pro-resolving effects. **a.** Visual representation of the analysed L3-L6 area of the spinal cord. This overview was generated using BioRender **b.** Quantification of phosphorylated p38 protein expression in the L3-L6 area of the spinal cord normalized on b-Actin (Western Blot) **c.** Quantification of c-Fos+ cells in the spinal cord and exemplary pictures of IF stained slides **d.** UMAP plot of spinal cord cells isolated from CIA ± RαMH and donut chart (% of live cells) of the different cell clusters analysed by spectral flow cytometry **e.** CD86+ inflammatory microglia **f.** TMEM119+ homeostatic microglia **g.** Clinical arthritis score shown as paw thickness (mm) of CIA mice with and without microglia depletion (25-30 dpi) ± RαMH at peak of disease. Data are expressed as the mean ± sd. Statistical difference was determined by and Student’s t-test (b,c) One-way ANOVA (e,f) and One-way ANOVA of area under the curve (g). *p < 0.05; **p < 0.01; ***p < 0.001; ****p < 0.0001.

### Oral H3R agonist treatment influences microglia function in the spinal cord

Our finding that microglia are essential for the intestinal histamine-induced resolution of arthritis prompted us to analyze their transcriptomes in CIA mice following oral H3R agonist treatment. Therefore, CD11b^+^ spinal cord microglia were isolated from CIA mice after oral H3R treatment and processed for bulk RNA sequencing (**Fig. 5a).** Microglia from H3R agonist treated CIA mice showed a significantly altered gene expression profile, predominantly characterized by an increase of immune modulatory genes such as *Fmr1nb, C4b, Spp1 and CD72 (61-64)* (**Fig. 5b**). Gene set enrichment analysis (GSEA) of hallmark genes revealed an overall anti-inflammatory phenotype. Inflammatory pathways such as IL-6, JAK/STAT3 and TNFα signaling were downregulated whereas anti-inflammatory pathways such as oxidative phosphorylation were upregulated (**Fig. 5c**). Further, analysis of the microglial microenvironment by untargeted metabolomics in spinal cords of H3R treated CIA mice identified a significantly changed secreted metabolite pattern with increased thiamine, acetylcholine and gamma-aminobutyric acid (GABA) in lumbar spinal cord tissue supernatants (**Fig. 5d**). Of note, identical spinal cord metabolites were previously linked to attenuated clinical scores in RA(65, 66) by reducing *p38* phosphorylation in the CNS(15, 67). Metabolite set enrichment analysis (MSEA) of identified spinal cord metabolites revealed the aspartate and purine metabolism as well as arginine biosynthesis amongst the highest enriched metabolites (**Fig. 5e)**. L-arginine was previously shown to inhibit arthritis and associated inflammatory bone loss in mice (68). In addition, RA patients exhibit lower purine metabolism activity (69), and a shortage of aspartate was shown to fuel synovial inflammation in RA(70). Together, these data identify oral histamine as a pro-resolving regulator of microglial gene expression and spinal cord metabolites.

**Figure 5:**
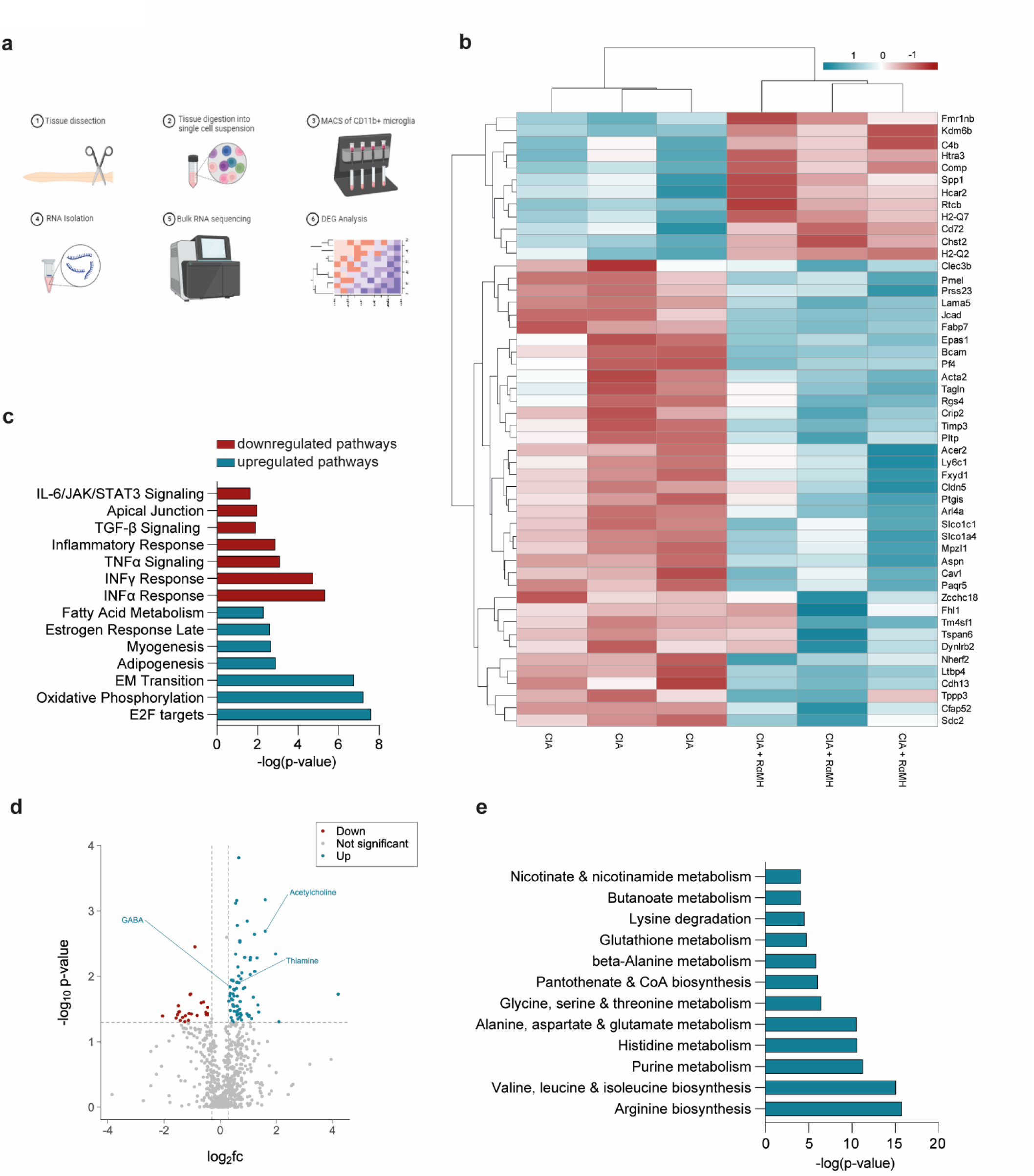
Oral H3R agonist treatment reverses the pro-inflammatory environment of the CNS. **a.** Layout of the RNAseq experimental layout. This overview was generated using BioRender **b.** Heatmap of the top 50 regulated genes in CD11b+ cells of the spinal cord (SC) in CIA mice ± RαMH **c.** Gene set enrichment of top 7 up- and downregulated hallmark pathways in the SC **d.** Volcano plot of untargeted metabolomics data of SC tissue supernatant of CIA mice ± RαMH. Cut-off were p-value < 0.05 and -0.3 < log2fc < 0.3 **e.** Most upregulated metabolic pathways identified using MetaboAnalyst.

### H3R signaling reduces the influx of cells to the joints

Microglia sense neuronal activity and can directly modulate their functions (71). In RA models, sciatic nerve branches were shown to control vascular leakage in arthritic paws (72-74). Therefore, plantar nerve fibers were isolated from the peak of activity of CIA following oral H3R agonist treatment. Sort-purified CD11b^-^ nerve cells were analyzed by bulk RNAseq (**Fig. 6a)**. The genes most differentially expressed included representatives of several signaling pathways critical for regulating cell and tight junction organization as well as GABA receptor activation and neurovascular coupling as indicated by enriched pathway analysis (**Fig. 6b)**. Further analysis of upstream regulators for neurovascular coupling signaling identified nuclear receptor (NR) NR4a3 and Mef2C as most significantly upregulated, both described to be involved in vascular biology and microglial inflammatory responses (75, 76) (**Fig. 6c, Supplementary Figure 7a, b)**. The Smoothelin-Like Protein 1 (SMTNL1) upstream regulator was most reduced in plantar nerve CD11b^-^ cells, similar to what was found in K/BxN mice following denervation of the sciatic nerve (74) (**Fig. 6c)**.

**Figure 6:**
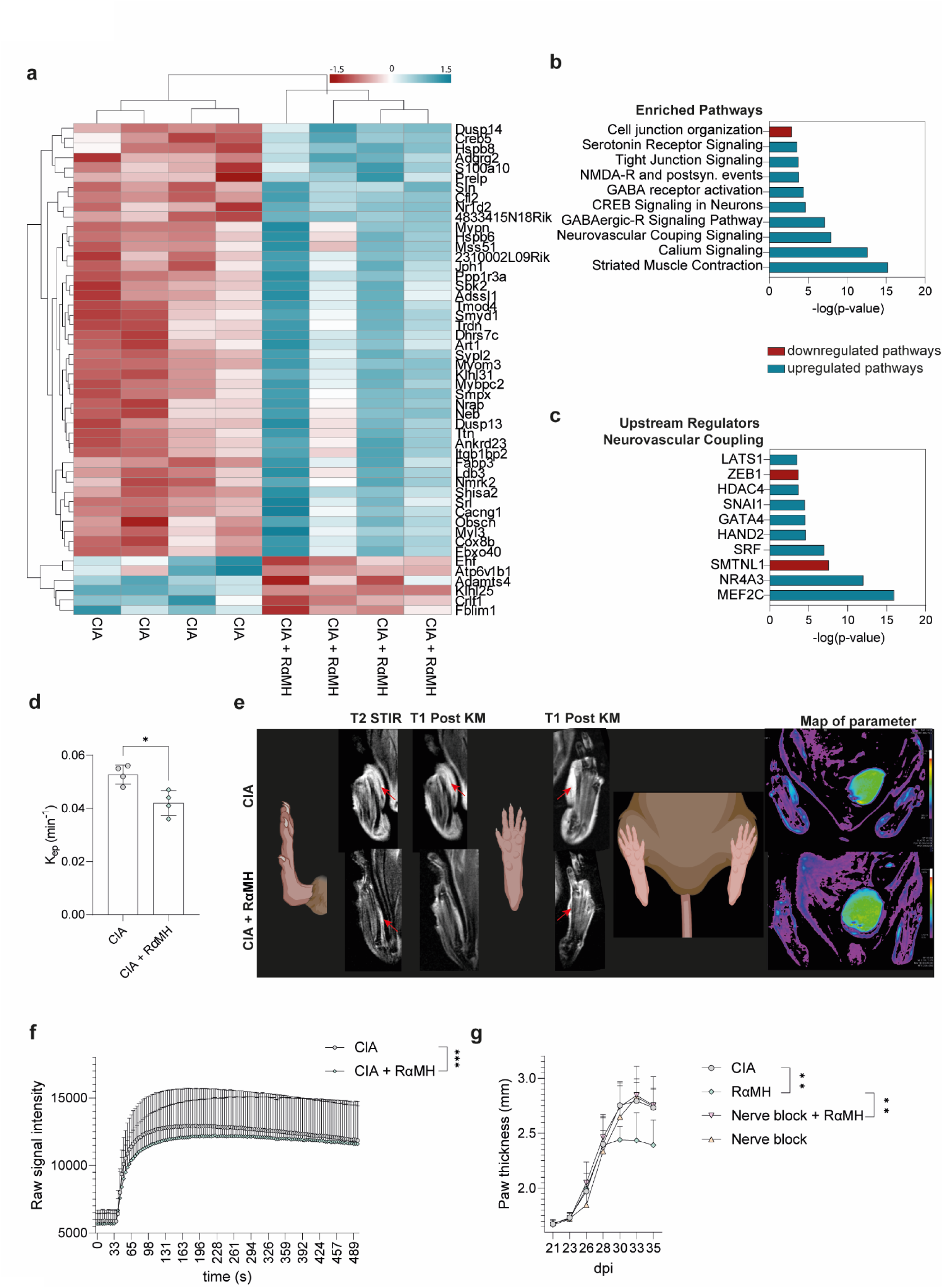
H3R signaling reduces the influx of cells to the synovium. **a.** Heatmap of the top 50 regulated genes in CD11b-cells of the Nervus plantaris (N.p.) in CIA mice ± RαMH **b.** Most enriched pathways identified by IPA analysis **c.** Top 10 upstream regulators of the enriched neurovascular coupling pathway (IPA) **d.** K_ep_ (Transfer constant) in the paws of CIA mice ± RαMH on third treatment day. **e.** Exemplary MRI pictures of T2 STIR, T1 POST KM, T1 POST KM and Map of parameter of CIA mice ± RαMH **f.** Raw signal intensity of DCE measurement over time of CIA mice ± RαMH **g.** Clinical arthritis score shown as paw thickness (mm) of CIA mice with/without QX-314 & bupivacaine induced nerve blockage ± RαMH (n=5) at peak of disease. Data are expressed as the mean ± sd. Statistical difference was determined by Student’s t-test (d), Student’s t-test of area under the curve (AUC) (f) and One-way ANOVA of AUC (g). *p < 0.05; **p < 0.01; ***p < 0.001

To analyze actual changes in vasoconstriction and vasodilation in CIA paws, *in vivo* magnetic resonance imaging (MRI) was performed at 28 and 31 dpi, before and after oral H3R agonist treatment, respectively. Reduced K_ep_ values, indicative of reduced vascular leakage, were found in H3R agonist-treated mice (**Fig. 6d)** along with overall lower raw signal intensity after contrast agent application (**Fig. 6e, f)**. Furthermore, the tendency to lower inflammatory area and paw thickness could be identified by MRT as soon as the last H3R agonist intervention administration (**Supplementary Fig. 7c, d)**. To investigate the dependence between the microglia changes in the spinal cord and the vascular leakage in the joints, we used a pharmacological approach to temporarily shut down nerve activity (73). QX-314, a membrane-impermeable lidocaine derivative that must enter a cell to block sodium channel conductance was locally injected in combination with bupivacaine in the footpad minutes before H3RA treatment at peak activity days of CIA. QX-314 treatment did not directly affect paw thickness in CIA mice, but abrogated the anti-inflammatory effects following oral H3R administration (**Fig. 6g)**. Together, this data suggest that local intestinal H3R activation restores vascular leakage in the inflamed joints by affecting centrally controlled local nerve innervation.

### Propionate supplementation increases local histamine levels in RA and MS patients

To translate our findings to human disease we utilized samples from two human studies, where RA and MS patients were supplemented with the SCFA propionate over the time course of several weeks. The publication of the initial clinical studies showed the beneficial effect of C3 or high fiber supplementation on disease activity of the patients (77-80). To elucidate if this effect of propionate is due to an increase in local histamine levels in the gut in the human setting, we analyzed stool samples from these patients at baseline and after 28 days (RA patients) or 90 days (MS patients) of C3 treatment. Interestingly, C3 supplementation over the course of 28 and 90 days significantly increased histamine levels in the stool samples of the patients (**Fig. 7a, b**). To further elucidate the role of microbiota-derived histamine in the setting of we reanalyzed RNAseq data from the *ex vivo* gut explant model utilized by Duscha et al (77), where the gut explants were treated with the microbiota of patients before and after C3 supplementation. Here, we looked at genes that are associated with H1R and H2R activation. Interestingly, H1R and H2R associated genes (histamine response network) were downregulated after C3 treatment (**Fig 7c**), although local histamine levels were significantly increased. This observation supports our hypothesis that this low-level histamine derived from microbiota acts via H3R activation and does not induce allergy or intolerance-associated reactions, mediated by H1R or H2R activation. Furthermore, histological analysis of H3R expression in ileal biopsies of RA patients and healthy controls revealed low levels of H3R expression in the healthy controls, consistent with previously published data (45). In RA patients, however, H3R expression in ileal tissues was strongly increased (**Fig 7d, e**). Taken together, these data suggest that our results in preclinical models can be directly transferred to human patients.

**Figure 7:**
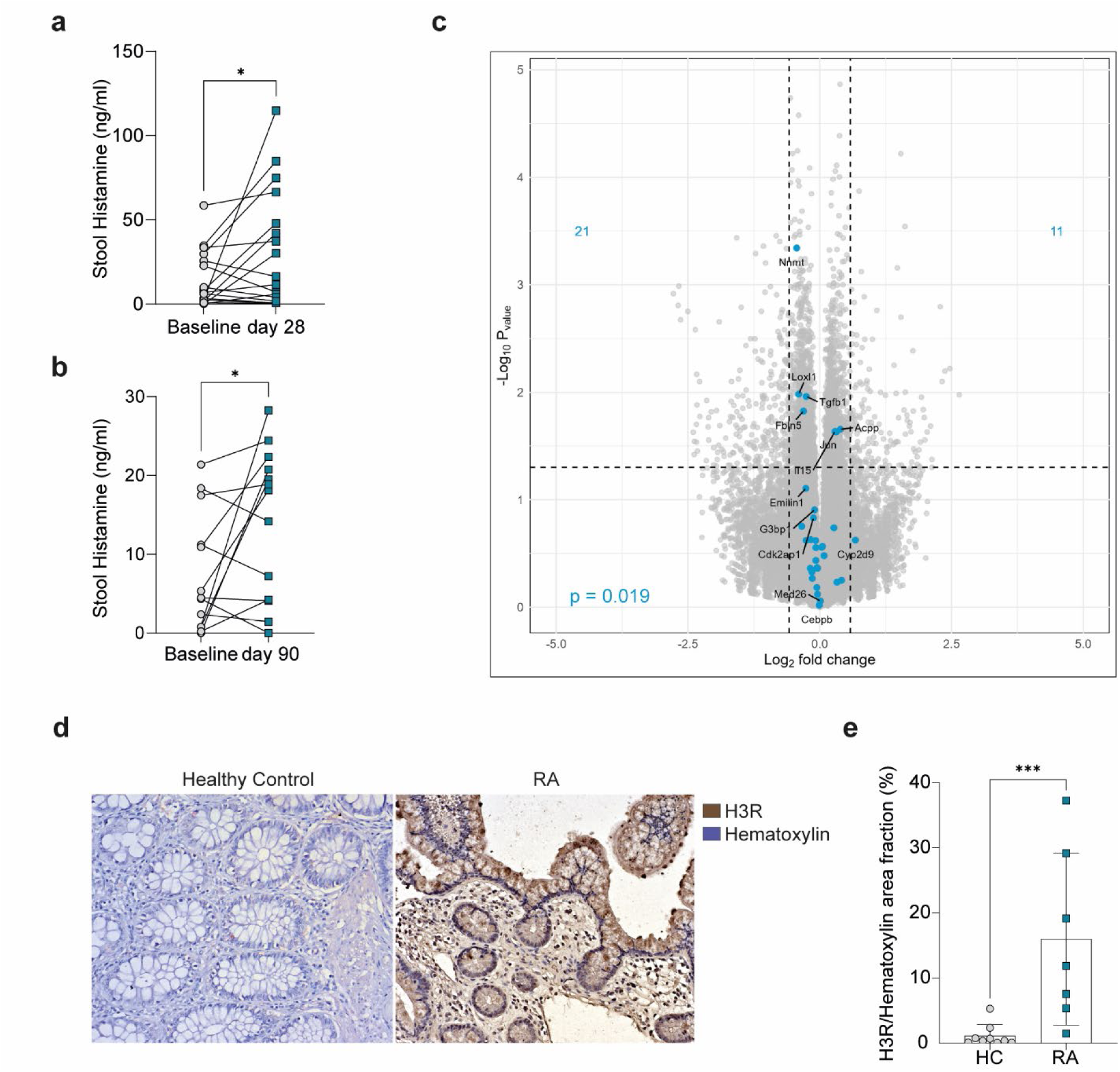
Propionate supplementation increases local histamine levels in RA and MS patients. **a.** Histamine levels in stool extracts from RA patients before and after 28 days of propionate supplementation **b.** Histamine levels in stool extracts from MS patients before and after 90 days of propionate supplementation **c.** Volcano Plot of RNAseq data from *ex vivo* colon culture infused with stool samples from patients before and after 90 days of propionate supplementation. Overlay with pathway genes of H1R and H2R activation (GSEA pathway POS HISTAMINE RESPONSE NETWORK) **d.** Exemplary pictures of histology staining of ileum biopsies from healthy controls (HC) or RA patients stained for H3R. **e.** Quantification of histological H3R staining of ileum biopsies from patients and HC. Data are expressed as the mean ± sd. Statistical difference was determined by Chi-squared test (a, b) and, Student’s t-test (e). *p < 0.05; ***p < 0.001.

## Discussion

The concept that the nervous system senses environmental stimuli and transmits these signals to immune cells to maintain tissue homeostasis is well-established (81). However, during chronic arthritic inflammation, it has been shown that the nervous system can exert both, pro-inflammatory or anti-inflammatory functions. For example, non-invasive electrical vagus nerve stimulation was shown effective to attenuate arthritis in CIA mice and RA patients (82, 83). On the other hand, denervation of the sciatic nerve protected mice from K/BxN serum transfer arthritis and individuals who suffer paralysis on one side of the body developed arthritis only on the neurologically unaffected contralateral side (74, 84).

So far, irrespective of the clinical effects, the regulation of arthritis by the CNS has only been investigated in a two-dimensional approach between the CNS and the joints. Here we identified spinal cord microglia as an essential gut-joint interface responsible for transmitting the anti-inflammatory pro-resolving effect of microbiota-derived histamine to the joints, thereby extending the neuro-immunomodulatory concept in RA to a third dimension, the gut.

Building on seminal work from both the Firestein (15-17) and Straub (85-87) who postulated the CNS as a new potential target for the treatment of rheumatic diseases, we experimentally confirmed their previous observations of reduced c-Fos protein expression and p38 phosphorylation locally in the spinal cord after i.th. application of p38 or TNFα inhibitors, simply by oral histamine supplementation. That is relevant because peripheral inflammation increases enhances the percentage of neurons with p38 activation in the spinal cords of mice (88).

By involving higher structures of the brainstem, the CNS effectively controls systemic immune responses broadly via the hypothalamic-pituitary axis (HPA) and the release of anti-inflammatory glucocorticoids by the adrenal cortex. These effects stand in contrast to the somatic sensory afferent fibres in the arthritic paws, which are under the local control of the spinal cord and send signals back to the periphery through so-called antidromic action potentials of the dorsal root reflex (DRR) (89, 90). This reflex is mainly pro-inflammatory through the recruitment of inflammatory cells and by promoting vasodilation and vessel leakiness as somatic denervation in mice was shown to protect from inflammatory arthritis (90). We could show by *in vivo* magnetic resonance imaging (MRI) that shortly after oral H3R agonist treatment vessel leakiness in the inflamed paws of CIA mice was likewise reduced and by temporarily shutting down nerve activity, the pro-resolving effect of histamine was lost.

The autonomic nervous system is part of the CNS that regulates involuntary body functions and is considered essential for the control of the regional homeostasis at the level of individual organs. The autonomic nervous system comprises two main branches: the sympathetic and parasympathetic. Parasympathetic stimulation was shown to suppress peripheral activation via the so-called cholinergic reflex that is triggered by the major parasympathetic neurotransmitter acetylcholine (ACh) (91, 92). Interestingly, ACh was among the most upregulated metabolites identified in spinal cord tissues following oral H3R agonist treatment in the current study, which is in accordance with previous reports showing H3R-dependent ACh release by histaminergic neurons (93). In preclinical RA models, ACh that was shown to effectively attenuate arthritis and genetic knockdown of the alpha7 nicotinic acetylcholine receptor (α7nAChR) increased CIA severity in mice (17, 66, 94, 95).

Next to ACh, we found GABA as significantly increased in spinal cord tissues following oral H3R treatment. GABA is a non-proteinogenic amino acid that is widely present in microorganisms, plants, and vertebrates and known to act as an inhibitory neurotransmitter in the CNS. Cadherin-13 (CDH13) was shown to be acritical regulator of GABAergic modulation (96) and was found upregulated amongst most significant genes in sort purified microglia following oral H3R agonist treatment. GABA agonist treatment effectively improved clinical signs in a rat model of chronic inflammatory pain (97). The role of GABA signalling in arthritis has also been investigated showing that the GABAergic system counteracting both the development and progression of RA (98), potentially by dampening T- and antigen-presenting cells (APC) activity (99). Notwithstanding the fact that GABA levels have been shown to be lower in people with RA compared to healthy individuals (100), the role of the GABAergic system in RA is complex and further research is needed to unravel its functions.

Acute peripheral inflammation was shown to increase glutamate in the spinal cord (101) as seen in the spinal cords of CIA mice following H3R agonist treatment. Glutamate, together with ACh and GABA, was described as potent modulator of the microglia phenotype, shifting them to an anti-inflammatory homeostatic phenotype (102, 103) that we mimic with the microglia phenotype after H3R agonist treatment. As shown by two independent studies on CIA-, or TNFα-transgenic arthritis mouse models and on post-mortem brain tissue from people with RA, microglia per se exhibit an inflammatory transcriptomic phenotype during active disease (104, 105). We identified myocyte-specific enhancer factor 2C (Mef2C) as strongest induced upstream regulator in sorted nerve cells following H3R agonist treatment and Mef2C was shown to restrain microglial inflammatory response (75). Pharmacological silencing of primary afferents in arthritic joints prevented microglial activation (73) and depletion of activated microglia in the EAE model delayed disease onset while microglia-specific deletion of the non-canonical NF-kappaB-inducing kinase (NIK) impaired EAE disease progression (57, 58). Together with our own results these findings suggest a central role of microglia in the gut-specific H3R signalling-mediated resolution of inflammation.

We propose that microbiota-derived histamine in the gut stimulates the enteric nervous system via H3R signalling (48). This tissue-specific response to histamine triggered the restoration of anti-inflammatory homeostatic microglia in the spinal cord where pro-inflammatory microglia were found to be enriched during pre-clinical arthritic models and RA patients (104, 105). Both the transcription profiles of the microglia and the secreted metabolites in the spinal cord create a local anti-inflammatory environment that causes a reduction in vasodilatation and an egress of inflammatory cells from arthritic paws via efferent nerve fibres. We also found that disrupting this gut-CNS-joint axis either by depleting microglia in the spinal cord or by blocking nerve fibres in the paws abolished the pro-resolving effect of histamine. In addition, high-fibre diet or direct SCFA-propionate supplementation increases local histamine concentrations in the gut of RA and MS patients thereby supporting resolution of inflammation.

## Methods

### Mice

Five to six week old WT C57BL/6N (Charles River) and DBA/1J mice (Janvier) were acclimated for 1 week, followed by a 3 week co-housing period before starting the experiments. All mice were maintained under specific pathogen-free conditions at the Präklinisches Experimentelles Tierzentrum (PETZ) Erlangen, Germany and approved by the local ethics authorities of the Regierung of Unterfranken (#55.2-2532-2-424, #55.2-2532-2-1674, #55.2.2-2532-2-1180). Supplementation of sodium propionate (Sigma-Aldrich) was done in the drinking water at a final concentration of 150 mM and changed every 3 days. The animals received water w/wo C3 and standard chow (Ssniff Spezialdiäten GmbH) ad libitum.

### Collagen-induced arthritis (CIA)

CIA was induced in 8-week-old female DBA/1J mice by subcutaneous injection at the base of the tail with 100 µl of 0.25 mg chicken type II collagen (CII; Chondrex) in complete Freund adjuvant (CFA; Difco Laboratory) containing 5 mg/ml killed Mycobacterium tuberculosis (H37Ra). Mice were re-challenged after 21 days intradermal immunization in the base of the tail with 100 µl of 0.25 mg chicken type II collagen (CII; Chondrex) in incomplete Freund adjuvant. The paws were evaluated for joint swelling three times per week. Each paw was individually measured for paw thickness using a caliper.

### Supplementation of Sodium Propionate in the drinking water

Mice were supplemented with 150 mM Sodium Propionate (C3, Sigma) in the drinking water. Drinking water was changed every two days to ensure consistent concentration and quality of the solution. Naïve mice were treated for 3 weeks as donors for fecal microbiota transfers (FMT), CIA mice were treated starting from peak of disease severity (30 dpi).

### Fecal microbiota transfer (FMT)

Donor mice were treated for at least three weeks with 150 mM sodium propionate (C3) in the drinking water. For transfer into five recipient mice, two donor mice were sacrificed, and their cecum contents were mixed with 2.5 ml 1 x PBS (for transfer) or H20 (for FPLC or untargeted metabolomics). The mixture was filtered through a 100 μm cell strainer. Then, 250 μl of the filtered solution was transferred into the donor mice by oral gavage. To guarantee the stability of the transfer, the procedure was repeated after two days. To assess the effect of soluble components or bacterial and cellular matters on ongoing CIA in mice the stool mixture was centrifuged at 3000 g. Supernatant fraction was aspirated and pellet was resuspended in PBS before transfer into mice with CIA by oral gavage.

### Culture and transfer of E. Coli strains

Frozen E. Coli glycerol stocks (kindly provided by Cezmi Akdis, Davos) were inoculated in 5 ml LB Medium (Roth). Bacteria were incubated overnight in a bacteria shaker at 37 °C, and 300 rpm. The next day the OD450nm was measured every hour until reaching the exponential phase. DBA1/J mice were treated with E. Coli B21 ± HDC by oral gavage. Each mouse received 1x108 per 250 µl dose reconstituted in PBS. Control mice received 250 µl PBS by oral gavage. Treatment was performed twice at days 30 and 32 post immunization.

### Fraction-generation by Fast protein liquid chromatography (FPLC)

Supernatant generated in paragraph 6.4 was further separated into 5 different size-specific fractions. In a first step sample was centrifuged at 100.000 g for 60 min. Supernatant was transferred on a VivaSpin 6 Column with a cut-off of 3 kDa and centrifuged for about 40-60 min until the remaining supernatant was 250 µl. Flow-through was collected as Fraction 1 for further transfer. Remaining supernatant was collected for separation into 4 further fractions by FPLC in cooperation with Xiang Wei from the Molecular Neurology Department of the UK Erlangen. Fractions were aliquoted and stored on -80 °C until transfer to mice. Supernatant fractions were thawed shortly before transfer into mice at peak of CIA by oral gavage. Mice were treated for 5 days with 250 µl of the corresponding fraction or PBS as a control

### Oral treatment with Histamine and Histamine-receptor agonists

Histamine and Histamine receptor agonists were dissolved in PBS to a stock-concentration of 0,5 mM and stored at -20 °C. Histamine treatment concentrations were carefully selected based on histamine concentrations in the stool of RA and MS patients after C3 treatment. Mice at peak of CIA were treated with 250 µl Histamine, H1R-Agonist, H2R-Agonist, H4R-Agonist (900 nM) [equipotent to histamine] or H3R-Agonist (60 nM) [15x more potent than histamine] by oral gavage for 3 days.

**Table 1:**
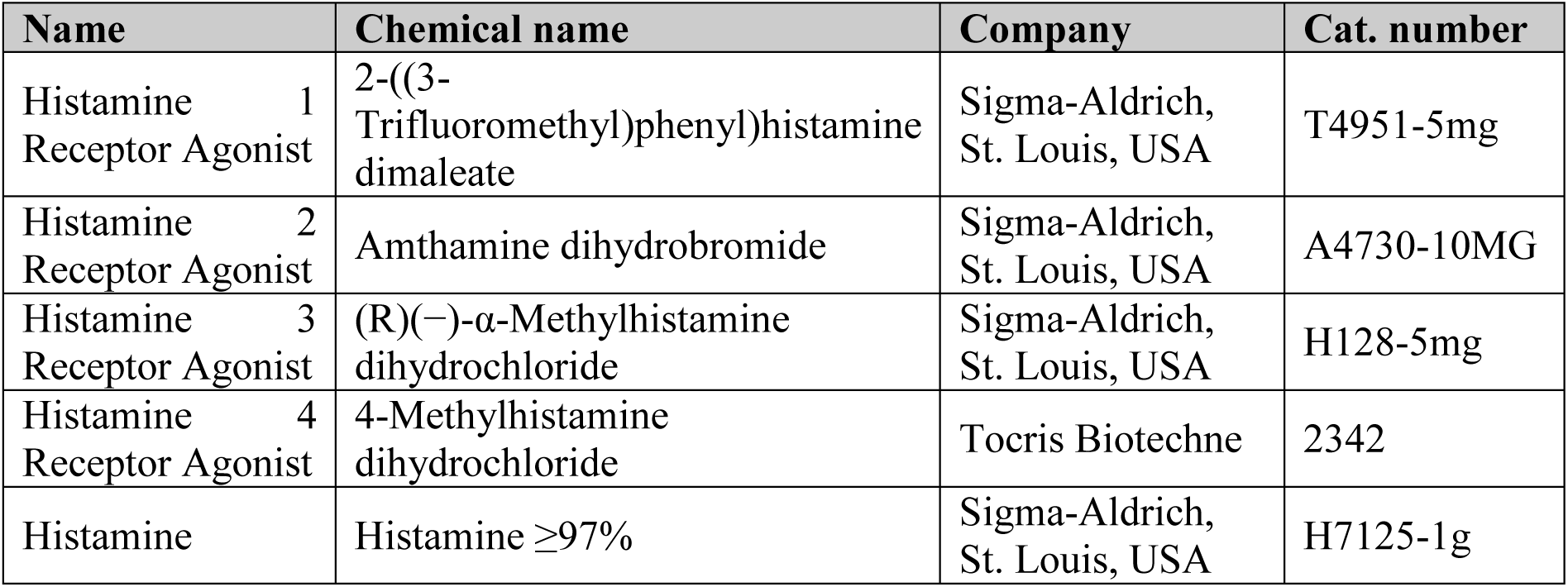
Used Histamine Receptor Agonists.

### Microglia depletion

Microglia were depleted using the CSF1R antagonist PLX5622. Mice received daily i.p. injections with 50 mg/kg PLX5622 (reconstituted in DMSO) or DMSO as control over 5 days.

### Blockage of primary afferents in the paws

The membrane impenetrable lidocaine sodium channel blocker QX-314 (2%, Sigma, 552233) was co-administered daily with bupivacaine (10 µg, Sigma, B5274) intra-plantarily in the hind paws for three days in combination with oral H3R agonist treatment in CIA mice at peak of disease days (28 – 30 dpi).

### EAE

EAE was induced in 8-12-week female C57Bl/6J mice using 150 μg of MOG35–55 (Genemed Synthesis, 110582) mixed with freshly prepared complete Freund’s Adjuvant (using 20 ml Incomplete Freund’s Adjuvant (BD Biosciences, #BD263910) mixed with 100 mg Myobacterium tuberculosis H37Ra (BD Biosciences, #231141)) at a ratio of 1:1 (v/v at a concentration of 5 mg/ml). All mice received two subcutaneous injections of 100 μl each of the MOG35-55/CFA mix. All mice then received a single intraperitoneal injection of 200 ng pertussis toxin (List Biological Laboratories, #180) at a concentration in 200 μl of PBS. Mice received a second pertussis toxin injection at the same concentration two days after EAE induction. Mice were monitored and scored daily thereafter. EAE clinical scores were defined as follows: 0, no signs; 1, fully limp tail; 2, hindlimb weakness; 3, hindlimb paralysis; 4, forelimb paralysis; 5, moribund.

### Tissue isolation

#### Spleen/mLN/pLN

Tissue was isolated, minced and mashed through a pre-wetted 70 μm cell strainer into 5 ml PBS. The cell suspension was centrifuged at 1500 rounds per minute (rpm) for 5 minutes (min), the supernatant was discarded and the pellet was resuspended in 3 ml RBC lysis buffer and incubated for 5 min. The RBC lysis was stopped by adding 10 ml FACS buffer, the samples were centrifuged at 1500 rpm for 5 min and the supernatant was discarded. The pellet was resuspended in 5 ml FACS buffer and 100 μl of the suspension was transferred into a 96 well U-bottom plate for counting. According to the count each sample was adjusted to a concentration of 107 cells/ml in FACS buffer. For the cytokine-staining isolated cells were restimulated. Therefore, 10^6^ cells were seeded into a V-bottom 96 well plate in 1X ReMed and incubated for 4 h at 37°C. All samples were washed with PBS, followed by viability staining for 30 min at 4°C in the dark with the Aqua NIR Fixable Viability Dye. After washing with FACS buffer the cells were incubated with a mix of extracellular binding antibodies (diluted in FACS buffer) and incubated for 30 min in the dark at RT. For intracellular staining, cells were washed with FACS buffer and fixed by using the FoxP3 Fixation and Permeabilization Kit. After fixation the samples were washed two times with permeabilization buffer and then incubated with 100 μl of the intracellular antibodies, diluted in permeabilization buffer, for at least 1 h in the dark. The samples were washed with permeabilization buffer and then reconstituted in FACS buffer until sample acquisition. Samples were acquired with a Beckman Coulter Cytoflex S or Cytek Norhtern Lights.

### Spinal Cord and plantaris nerve

To isolate the cells from the PNS and CNS, mice were euthanized and perfused with cold 1× PBS. The nerves were isolated and collected on ice in 500µl digestion medium consisting of 0.15% Trypsin (Sigma-Aldrich, #T1426) diluted in medium and 0.3% (w/v) Collagenase II (Sigma-Aldrich, #11088858001). The tissue samples were mechanically diced in pieces using small microdissecting scissors and then shaken at 850 rpm for 10 minutes at 37 °C to facilitate enzymatic digestion. After trituration with a 1ml pipette tip, the samples were digested another 10 minutes on an orbital shaker at 850 rpm at 37 °C. The cell suspensions were then triturated with a 200µl pipette tip. Warm complete medium consisting of DMEM + GlutaMAX (Thermo Fisher Scientific Scientific, #61965026) supplemented with 10% FBS (Thermo Fisher Scientific Scientific, #10438026) and 1% penicillin/streptomycin (Thermo Fisher Scientific Scientific, #10500064) was added to stop digestion. The homogenized tissue was then mechanically dissociated using a 5-ml serological pipette and triturated through a 100-µm cell strainer (Fisher Scientific, #22363548) into a fresh 15-ml conical tube. Following this, the samples were centrifuged at 400g for 7 minutes at 4 °C. After discarding the supernatant, the remaining mononuclear cell pellet was resuspended in 200 µl of 1x PBS and prepared for further applications. CD11b+ microglia were isolated from the cell suspension using the CD11b Microbead Isolation kit (Miltenyi, #130-049-601) according to the manufacturer’s instructions.

### Flow Cytometry

#### mLN, pLN, Spleen, Paws

Organs were harvested, and single-cell suspensions were prepared. For flow cytometric analyses, cells were stained with antibodies to the following markers: Data was acquired on a Cytek Aurora Northern Lights (Cytek). Please refer to Supplementary Table 1 for further details on antibody clone, dilution factor, provider, target species and application (Suppl. Table 1). For dimensionality reduction, samples were down-sampled using the DownSample plugin (version 3.3.1). Clusters were first calculated with phenograph (version 2.4). T-SNE reduction was done with Flowjo (version 10.8.1). Then, clustering was done with FlowSOM (version 3.0.18) using the phenograph cluster number as input.

### Spinal Cord

For the characterization of CNS cells we used flow cytometric analyses. The LIVE/DEAD™ Fixable Aqua Dead Cell Stain Kit (Thermo Fisher Scientific, #L34957) was used to differentiate between dead and viable cells according to the manufacturer’s instructions. The cells were washed with cold 1x PBS and then incubated with Fc block (Purified Rat Anti-Mouse CD16/CD32; BD; #553141) in PBS for 10 minutes in the dark at room temperature. The cells were labelled with flow cytometry antibodies at 4°C in the dark for 30 minutes, diluted in FACS buffer (1x PBS, 2% FBS, 2 mM EDTA). After two washing steps in cold FACS buffer, the samples were resuspended in 1× PBS for acquisition. The following antibodies were used in the study: BV421-CD11b (Biolegend, #101235), BV480-CD11c (BD, #565627), BV570-Ly6C (Biolegend, #128029), BV605-CD68 (Biolegend, #137021), BV650-CD56 (BD, #748098), PE-eFlour610-CD140a (Thermo Fisher Scientific, #61140180), SuperBright780-MHCII (Thermo Fisher Scientific, #78532080), AF488-H3R (R&D, #FAB10200G), eFluor 450-CD3 (Thermo Fisher Scientific, #48003742), PE-Cy5-CD24 (Biolegend, #101811), PE-Cy7-CD31 (Thermo Fisher Scientific, #25031182), PerCP-eFlour710-CD86 (Thermo Fisher Scientific, #46086280), PE-B220 (BD, #561878), PE-Cy5.5-CD45 (Thermo Fisher Scientific, #35045180), APC-Cy7-Ly6G (Biolegend, #127623), AF700-O4 (R&D, #FAB1326N), BUV737-CD154 (BD, #741735), AF660-CD19 (Thermo Fisher Scientific, #606019380), APC/Fire810-CD4 (Biolegend, #100479), APC-ACSA2 (Miltenyi Biotech; #130-117-386). For data analysis, the OMIQ platform was used. In more detail, cells were gated according to previously described methods 43,49. For dimensionality reduction, cells were downsampled to an appropriate number per group. Opt-SNE was performed (maximum 1000 iterations, perplexity 30, theta 0.5, verbosity 25) or UMAP (15 neighbors, minimum distance 0.4, 200 Epochs), followed by PhenoGraph clustering based on Euclidean distance. When comparing two groups, a two-class unpaired approach using SAM (Significance Analysis of Microarrays) was performed, with a maximum of 100 permutations and a False Discovery Rate (FDR) cutoff of 0.1.

### µCT

µCT imaging was performed using the cone-beam Desktop Micro Computer Tomograph “µCT 40” by SCANCO Medical AG, Bruettisellen, Switzerland. The settings were optimized for calcified tissue visualization at 55 kVp with a current of 145 µA and 200 ms integration time for 500 projections/180°. For the segmentation of 3D-volumes, an isotropic voxel size of 8.4 µm and an evaluation script with adjusted grayscale thresholds of the operating system “Open VMS” by SCANCO Medical was used. Volume of interest tibia: The analysis of the bone structure was performed in the proximal metaphysis of the tibia, starting 0.43 mm from an anatomic landmark in the growth plate and extending 1.720 mm (200 tomograms) distally.

### Histology

Tibial bones were fixed in 4% formalin for 24 h and decalcified in EDTA (Sigma-Aldrich). Serial paraffin sections (2 μm) were stained for tartrate resistant acid phosphatase (TRAP) using a Leukocyte Acid Phosphatase Kit (Sigma) according to the manufacturer’s instructions.

Osteoclast numbers were quantified using a microscope (Carl Zeiss) equipped with a digital camera and an image analysis system for performing histomorphometry (Osteomeasure; OsteoMetrics). For Calcein labeling, mice were injected with 30 mg/kg body weight of green fluorescent Calcein (Sigma) 11 and 4 days before sacrifice. Undecalcified bones were embedded in methacrylate and 5 mm sections were cut. Unstained sections were used to measure fluorescent Calcein-labeled bone surfaces at a wavelength of 495 nm. Toluidine Blue staining was performed for quantification of osteoblasts and von Kossa staining for bone mineralization.

### SCFA measurement

Four to five replicates of frozen cecal samples (100 mg) or 50 µl of serum were weighed into a 2 ml polypropylene tube. The tubes were kept in a cool rack throughout the extraction. 33% HCl (50 µl for cecal contents or 5 µl for serum) was added and samples were vortexed for 1 min. One milliliter of diethyl ether was added, vortexed for 1 min, and centrifuged for 3 min at 4 °C. The organic phase was transferred into a 2 ml gas chromatography (GC) vial. For the calibration curve, 100 μl of SCFA calibration standards (Sigma) were dissolved in water to concentrations of 0, 0.5, 1, 5, and 10 mM and then subjected to the same extraction procedure as the samples. For GC mass spectrometric (GCMS) analysis, 1 μl of the sample (4–5 replicates) was injected with a split ratio of 20:1 on a Famewax, 30 m × 0.25 mm iD, 0.25 μm df capillary column (Restek, Bad Homburg). The GC-MS system consisted of GCMS QP2010Plus gas chromatograph/mass spectrometer coupled with an AOC20S autosampler and an AOC20i auto injector (Shimadzu, Kyoto, Japan). Injection temperature was 240 °C with the interface set at 230 °C and the ion source at 200 °C. Helium was used as carrier gas with constant flow rate of 1 ml/min. The column temperature program started with 40 °C and was ramped to 150 °C at a rate of 7 °C/min and then to 230 °C at a rate of 9 °C/min and finally held at 230 °C for 9 min. The total run time was 40 min. SCFA were identified based on the retention time of standard compounds and with the assistance of the NIST 08 mass spectral library. Full-scan mass spectra were recorded in the 25–150 m/z range (0.5 s/scan). Quantification was done by integration of the extracted ion chromatogram peaks for the following ion species: m/z 45 for acetate eluted at 7.8 min, m/z 74 for propionate eluted at 9.6 min, and m/z 60 for butyrate eluted at 11.5 min. GCMS solution software was used for data processing.

### 16s rRNA sequencing

To extract genomic DNA from stool samples, the Qiamp Fast DNA Stool extraction kit (Qiagen, Venlo, Netherlands) was used, following the manufacturer’s instruction. The NEBNext Q5 Host Start Hifi PCR Master Mix (NEB) was utilized for amplification of the V3– V4 region of the bacterial 16S rRNA gene. The purification of amplified fragments was done with AMPure XP Beads (Beckmann Coulter Genomics, Brea, California). This was combined and analyzed by a “2 × 300 bp paired-end” sequencing on a Illumina MiSeq machine. The Usearch10 MicrobiomeAnalyst package [10] was used to perform a quality control, OTU table generation and bioinformatics analysis [11]. Classification was done by utilizing the “Ribosomal database project” (RDP release 16).

### Untargeted metabolomics of intestinal supernatant, serum and spinal cord extracts

#### Sample preparation

In order to extract analytes, stool suspensions of feces (in H_2_O) were centrifuged at 16000 rpm and 4°C for 5 min using an Eppendorf 5427R centrifuge (Eppendorf; Hamburg, Germany). 100 µL of supernatant were mixed with 400 µL of ice-cold methanol containing recovery standards for evaluation of the quality of cell harvest and correction for variations. These standards were tridecanoic acid (25 µg/mL), DL-2-fluorophenylglycine (1.25 µg/mL), [2H6]-cholesterol (25 µg/mL), and DL-4-chlorophenylalanine (0.02 µg/mL). This was done for both HILIC and RP samples (see below). Before preparation for LC-MS analysis each sample was centrifuged at 16000 rpm and 4°C for 5 min using an Eppendorf 5427R centrifuge (Eppendorf; Hamburg, Germany). From each supernatant 350 µL were pipetted into two separate LC vials. 230 µL of each unknown sample were pooled to receive pooled QC samples. Of this pool 350 µL supernatant were pipetted into LC vials for each QC sample.

All samples were dried under a gentle stream of nitrogen at 30°C. Following to this, samples were reconstituted in 200 µL of eluent for HILIC analysis or eluent for RP analysis, to resemble conditions at the beginning of each chromatographic run (see next section for details). The eluents contained internal standards for evaluation of instrumental performance for each sample.

#### LC-Orbitrap-MS-Analysis

LC-MS analysis was performed as previously described (106). In brief, high-resolution mass spectrometry was done on an Exactive Focus Hybrid Quadrupole-Orbitrap mass spectrometer coupled to a Dionex Ultimate 3000 chromatographic system (both from Thermo Fisher Scientific, Dreieich, Germany). Chromatographic separation was performed via both HILIC and RP chromatography. For HILIC separation, an Acquity UPLC BEH Amide, 1.7 µm, 2.1 x 100 mm column was used, while for RP an Acquity UPLC BEH C18, 1.7 µm, 2.1 x 100 mm column was installed. Both columns were equipped with a 2.1 x 5 mm guard column, respectively (all columns from Waters, Eschborn, Germany). The applied mass spectrometric settings, LC gradients, run times, QC limits, and batch sequences were identical to those previously shown by Gessner et al (106).

#### Data analysis

Data was analyzed with Compound Discoverer 3 (Thermo Fisher Scientific, Dreieich, Germany). Features were grouped and identified with a mass tolerance of 5 ppm and a retention time tolerance of 0.2 min. Differential analysis was performed using the log-transformed values for the peak area of a feature. P-values of ≤ 0.05 were considered as statistically significant.

### Metabolite set enrichment analysis

Metabolite set enrichment analysis was performed using the MetaboAnalyst webtool (107). Only significantly increased level 1 annotated metabolites were used for this analysis.

### Measurement of serum cytokines and chemokines (Legendplex)

Serum cytokines were measured with the LEGENDplex™ Mouse Proinflammatory Chemokine Panel (#740007, Biolegend) and LEGENDplex™ MU Th Cytokine Panel (12-plex) (#741044, Biolegend) following the manufacturer’s instructions.

### Ex vivo gut organ culture and c-FOS staining

Fabrication of the gut organ culture device and gut organ culture experiments were performed as previously described (46, 108). Briefly, intact whole colons were dissected sterilely from 14d old C57BL/6 mouse littermates reared under SPF conditions. The solid lumen content was gently flushed, and the gut fragment was threaded and fixed over the luminal input and output ports of the gut organ culture device, using sterile surgical thread. The culture device was placed in a custom-made incubator that maintains a temperature of 37c, and tissue was maintained half-soaked in a constant flow of sterile medium (Iscove’s Modified Dulbecco’s Medium (IMDM) along with 20% Knockout serum, 2% B-27, 1% N-2, 1% L-glutamine, 1% non-essential amino acids, and 1% HEPES) using a syringe pump. Gut cultures were infused with Histamine H3RA agonist at 250nM into the gut lumen using a syringe pump (littermate tissues infused with sterile medium served as an internal control). After 1 hour, tissues were collected for whole mount staining. Tissues were fixed with cold (4c) 4% PFA for 1h. Tissues were then permeabilized at room-temperature using 0.5% triton for 2h and then transferred into blocking solution (0.1% triton, 5% BSA, and 10% donkey serum in PBS). Tissue were incubated with primary antibodies against beta-III-tubulin (pan-neuronal marker, Abcam, ab41489, 1:100) and c-Fos (Cell Signaling, cst2250s; 1:150), overnight at 4c. Tissues were then washed and incubated with secondary antibodies (Cy3 (rabbit) and Cy5 (chicken), Jackson, 1:100) either for 2h, and then washed and counterstained with DAPI (1:1000) for 15 min. Colons were imaged using confocal fluorescence microscope (Leica Stellaris) and cFos nuclear localization was calculated using ImageJ. Briefly, ImageJ macro calculated nuclear c-Fos mean fluorescence intensity (MFI) in beta-III-tubulin+DAPI+ regions. The MFI values in individual nuclei were normalized to the average MFI in the internal control tissues, and statistical analyses were performed using GraphPad Prism 9 software.

### RNAseq of SC and N. plantaris

RNA from CD11b+ and CD11b+ tissue suspension of SC and N. plantaris was isolated using the QiaGEN RNeasy Micro Kit following manufacturer’s instructions. BulkRNAseq was performed by the Core Unit Next Generation Sequencing of the University Clinic Erlangen.

### IPA

RNAseq data were further analyzed for enriched pathways and upstream regulators with the use of QIAGEN IPA (QIAGEN Inc., https://digitalinsights.qiagen.com/IPA) (109).

### RNAseq of ileum

RNA from Ileal tissue samples from CIA mice ± oral RαMH treatment was isolated using the QiaGen RNeasy Mini Kit following the manufacturer’s instructions. Novogene Sequencing – Europe (UK, Cambridge Sequencing Center) performed the Illumina RNA sequencing (RNAseq). In brief, sequencing libraries were generated using NEBNext® Ultra TM RNA Library Prep Kit for Illumina® (NEB, USA) and sequenced on an Illumina platform. Raw data (raw reads) of FASTQ format were processed through fastp. Mapping of the processed data to the reference genome Mus Musculus (GRCm38/mm10) was performed using the Spliced Transcripts Alignment to a Reference (STAR) software (110). FeatureCounts was used for the quantification of the mapped reads (111). Raw mapped reads were processed in R (Lucent Technologies) with DESeq2 (112), to determine differentially expressed genes and generate normalized read counts. Pathway enrichment analysis was performed using the free online platform DAVID (113).

### GSEA

Gene set enrichment analysis of RNAseq data was performed using the ClusterProfiler Package (Version 4.10.0) in R (114).

### WesternBlot

For SDS-Page and Western Blotting the NuPAGE system from ThermoFisher Scientific was used. Tissues were homogenized in RIPA buffer containing proteinase and phosphatase inhibitors. Protein extracts were separated on a NuPAGE™ 4-12% Bis-Tris Protein Gels, transferred on a PVDC membrane and stained with antibodies against p38 (Cell Signaling #9218S) and phosphorylated p38 (Cell Signaling, #9211S). An antibody against β-Actin was used as loading control.

### MRI imaging

In vivo imaging by MRI of the hind paws was performed at days 28 (before oral RαMH treatment) and 31 after CIA immunization (after 3 days of RαMH treatment) with a preclinical 7T MRI (ClinScan 70/30, Bruker BioSpin, Ettlingen, Germany) using a volume resonator: RF RES 300 1H 075/040 QSN TR. The imaging protocol included T1-weighted spin echo sequences, a short tau inversion recovery (STIR) sequence and a Dynamic Contrast Enhanced (DCE) measurement using a fast low-angle shot (FLASH) sequence (Table 2). During DCE-MRI the mice received an intravenous bolus injection of a low molecular weight gadolinium chelate agent (0.15 mmol/kg Gadobutrol, Gadovist, Bayer Vital GmbH, Leverkusen, Germany) over a time period of 10 s via a tail vein catheter.

**Table 2:**
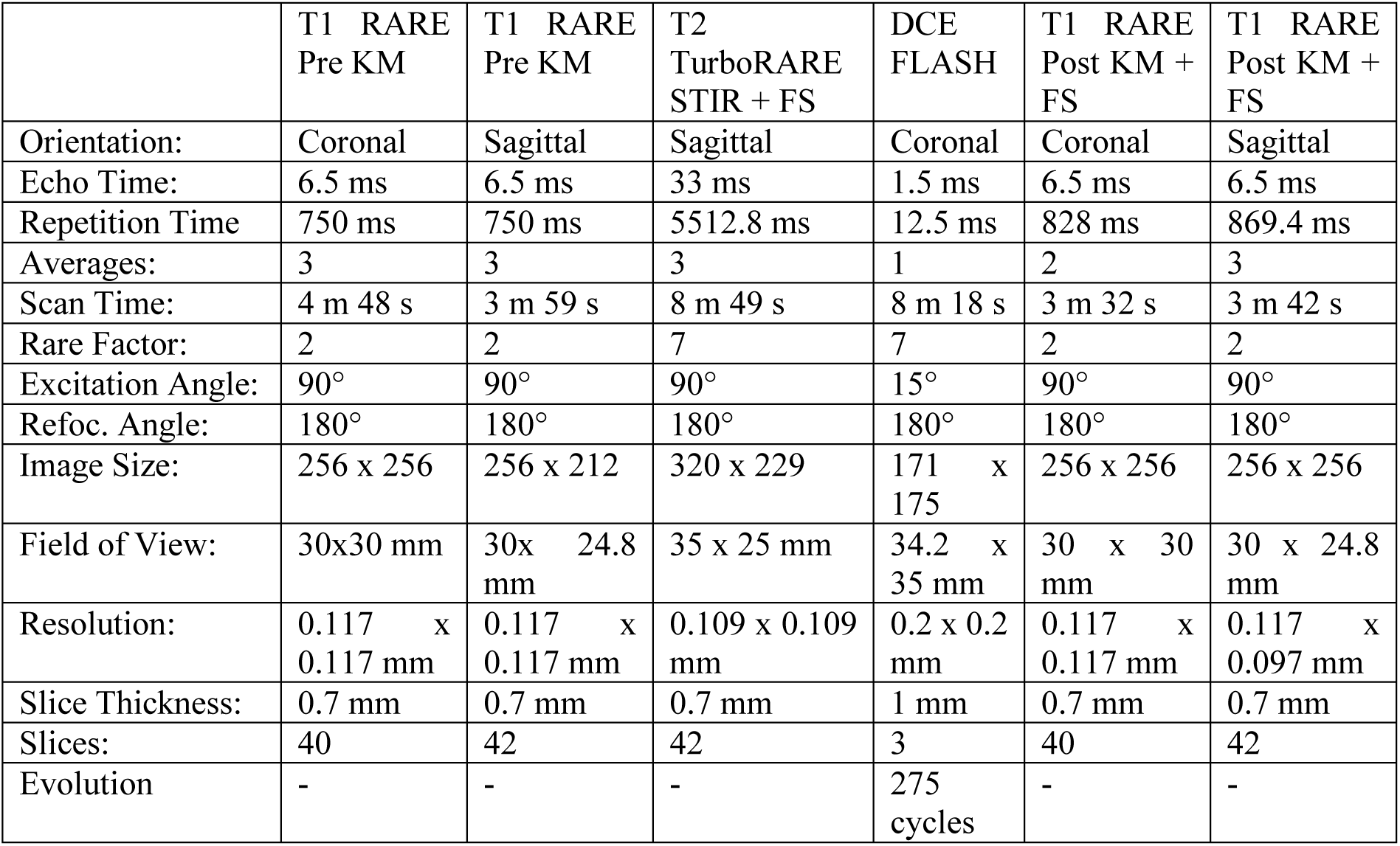
Imaging Protocols for the different MRI used sequences.

|  | T1 RARE<br>Pre KM | T1 RARE<br>Pre KM | T2<br>TurboRARE<br>STIR + FS | DCE<br>FLASH | T1 RARE<br>Post KM +<br>FS | T1 RARE<br>Post KM +<br>FS |
| --- | --- | --- | --- | --- | --- | --- |
| Orientation: | Coronal | Sagittal | Sagittal | Coronal | Coronal | Sagittal |
| Echo Time: | 6.5 ms | 6.5 ms | 33 ms | 1.5 ms | 6.5 ms | 6.5 ms |
| Repetition Time | 750 ms | 750 ms | 5512.8 ms | 12.5 ms | 828 ms | 869.4 ms |
| Averages: | 3 | 3 | 3 | 1 | 2 | 3 |
| Scan Time: | 4 m 48 s | 3 m 59 s | 8 m 49 s | 8 m 18 s | 3 m 32 s | 3 m 42 s |
| Rare Factor: | 2 | 2 | 7 | 7 | 2 | 2 |
| Excitation Angle: | 90° | 90° | 90° | 15° | 90° | 90° |
| Refoc. Angle: | 180° | 180° | 180° | 180° | 180° | 180° |
| Image Size: | 256 x 256 | 256 x 212 | 320 x 229 | 171 x<br>175 | 256 x 256 | 256 x 256 |
| Field of View: | 30x30 mm | 30x 24.8<br>mm | 35 x 25 mm | 34.2 x<br>35 mm | 30 x 30<br>mm | 30 x 24.8<br>mm |
| Resolution: | 0.117 x<br>0.117 mm | 0.117 x<br>0.117 mm | 0.109 x 0.109<br>mm | 0.2 x 0.2<br>mm | 0.117 x<br>0.117 mm | 0.117 x<br>0.097 mm |
| Slice Thickness: | 0.7 mm | 0.7 mm | 0.7 mm | 1 mm | 0.7 mm | 0.7 mm |
| Slices: | 40 | 42 | 42 | 3 | 40 | 42 |
| Evolution | - | - | - | 275<br>cycles | - | - |

### MRI analysis

The total volume each hind paw was segmented on T2 STIR images. The volume of edema (bound tissue water) was quantified by segmenting all voxels in the hind paws with signal intensity above 5500 a.u. on STIR using a dedicated threshold-based OsiriX (aycan OsiriX PRO v. 2.08.006, aycan Medical Systems, LLC) Plugin, the Chimaera software (Chimaera GmbH Erlangen, Germany). DCE data was analyzed by using Bruker ParaVision 360 V3.2 and Microsoft Excel (Microsoft Office Professional Plus 2019).

### Primary mouse astrocyte and microglia cultures and stimulation experiments

Brains of mice aged P0–P3 were dissected into PBS on ice. Brains of 6-8 mice were pooled, centrifuged at 500 × g for 10 min at 4 °C and resuspended in 0.25% Trypsin-EDTA (Thermo Fisher Scientific Scientific, #25200-072) at 37 °C for 10 min. DNase I (Thermo Fisher Scientific Scientific, #90083) was added at 1 mg/ml to the solution, and the brains were digested for 10 more minutes at 37 °C. Trypsin was neutralized by adding DMEM + GlutaMAX (Thermo Fisher Scientific Scientific, #61965026) supplemented with 10% FBS (Thermo Fisher Scientific Scientific, #10438026) and 1% penicillin/streptomycin (Thermo Fisher Scientific Scientific, #10500064), and cells were passed through a 70-µm cell strainer. Cells were centrifuged at 500 × g for 10 min at 4 °C, resuspended in DMEM + GlutaMAX with 10% FBS 1% penicillin/streptomycin and cultured in T-75 flasks (Sarstedt, #83.3911.002), pre-coated with 2 µg/ml Poly-L Lysine (PLL, Provitro, #0413) at 37 °C in a humidified incubator with 5% CO2 for 5–7 days until confluency was reached. Mixed glial cells were shaken for 30 min at 180 rpm, the supernatant was collected and the medium was changed, and then cells were shaken for at least 2 h at 220 rpm and the supernatant was collected and the medium was changed again. CD11b+ microglia were isolated from the collected supernatant using the CD11b Microbead Isolation kit (Miltenyi, #130-049-601) according to the manufacturer’s instruction. For stimulation experiments, astrocytes and microglia were detached using TrypLE (Thermo Fisher Scientific, #12604013) and seeded in PLL-coated 48-well plates (Sarstedt, #NC1787625) at a density of 150.000 cells per well.

### Human studies (ProDarMi/MS)

The RA C3 supplementation study was approved by the ethics committee of the Department of Medicine at the Friedrich-Alexander University Erlangen-Nürnberg (#431_20 B). The study protocol and Trial registration can be found in the German Clinical Trials Register (ID: DRKS00023985). The MS study was approved by the ethics committee of the Department of Medicine at the Ruhr-University Bochum (registration number 15-5351, 4493-12, 17-6235) and all study information has previously been published (77).

### Histamine ELISA of human serum and stool

For Histamine measurement in stool samples from MS and RA patients, stool extracts were isolated from stool samples using the Stool preparation system, filled with extraction buffer IDK® Amino Extract (#K7999, ImmunDiagnostik). Histamine was measured using the Histamin Stool ELISA Kit (#K8213, ImmunDiagnostik) using the manufacturer’s instructions and measured using a Tecan Sunrise Plate Reader.

### Data analysis and statistics

Statistical analyses were performed using Prism 9 software (GraphPad). Comparisons between two groups were performed using unpaired or paired, two-tailed, Student’s t test. Comparisons between more than two groups were performed using one-way ANOVA and Tukey’s or Dunnett’s multiple comparison test. Statistical significance: * p < 0.05; ** p < 0.01; *** p < 0.001; **** p < 0.0001. Details on the statistical analysis are listed in the figure legends.

## Data availability

All relevant data are available from the authors upon reasonable request. The source data underlying Figures 1-7 and Supplementary Figures 1-7 are provided as source data file. All data are available from the authors upon request.

## Supplementary Materials

Supplementary Figures 1-7.

Supplementary Table 1

## Acknowledgements

We thank the Optical Imaging Center Erlangen (OICE), Erlangen, Germany for their technical help with microscopy. We thank the Core Unit Next Generation Sequencing, Erlangen, Germany for their support. We thank Dr. Z. Winter and C. Engert from the Preclinical Imaging Platform Erlangen (PIPE) for their support with the MRI. We thank Prof. Sonnewald for his support. We thank the animal facility PETZ, Erlangen, Germany for supporting this project. We thank D. Weidner for her support with the µCT analysis. We thank B. Happich and N. Berndt from the Med3 Histology Lab for their support. We thank the Med 3 Studienambulanz for their support in collecting the study samples. We thank all members of our laboratories at the Medical Clinic 3, Erlangen, Germany for their support and helpful discussion. This study received funding from the Deutsche Forschungsgemeinschaft (DFG, German Research Foundation) via DFG-RU2886-Project A01 as well as DFG-CRC1181-Project-No. B07. This study was further funded by the Interdisciplinary Center for Clinical Research, Erlangen (IZKF) (project number P144) at the Universitätsklinikum Erlangen, Germany.

## Author contributions

G.S., V.T. and M.M.Z. designed the project, interpreted results and wrote the manuscript. K.D. and M.L. performed most of the work, analysed data, edited the manuscript and made figure panels. E.S., H.D., V.A., S.L., F.S., M.F., L.E. and L.L. acquired data and provided help with multiple experiments. L.A, T.R.L. and T.S. performed and analysed 16S RNA data. H.BM, H.R., Y.R. and N.Y. performed and analysed gut explant experiments. A.G. and R.V.T. performed and analysed untargeted metabolomics experiments. J.H. performed and analysed the SCFA measurements. D.M. and F.C. performed and analysed human H3R staining. F.B. and R.B. performed spinal cord histology. W.X. performed size exclusion chromatography. A.H. and C.A.A. provided human MS stool samples and HDC competent E.coli strains, respectively. T.B. performed and analysed MRT measurements. T.B. and K.S. wrote the animal licence approvals. M.M.Z supervised the project and provided funding. All the authors revised and approved the manuscript.

