## Supplementary Figures for "Gut-specific H3R signaling orchestrates microglia-dependent resolution of peripheral inflammation"

Sup. Fig. 1

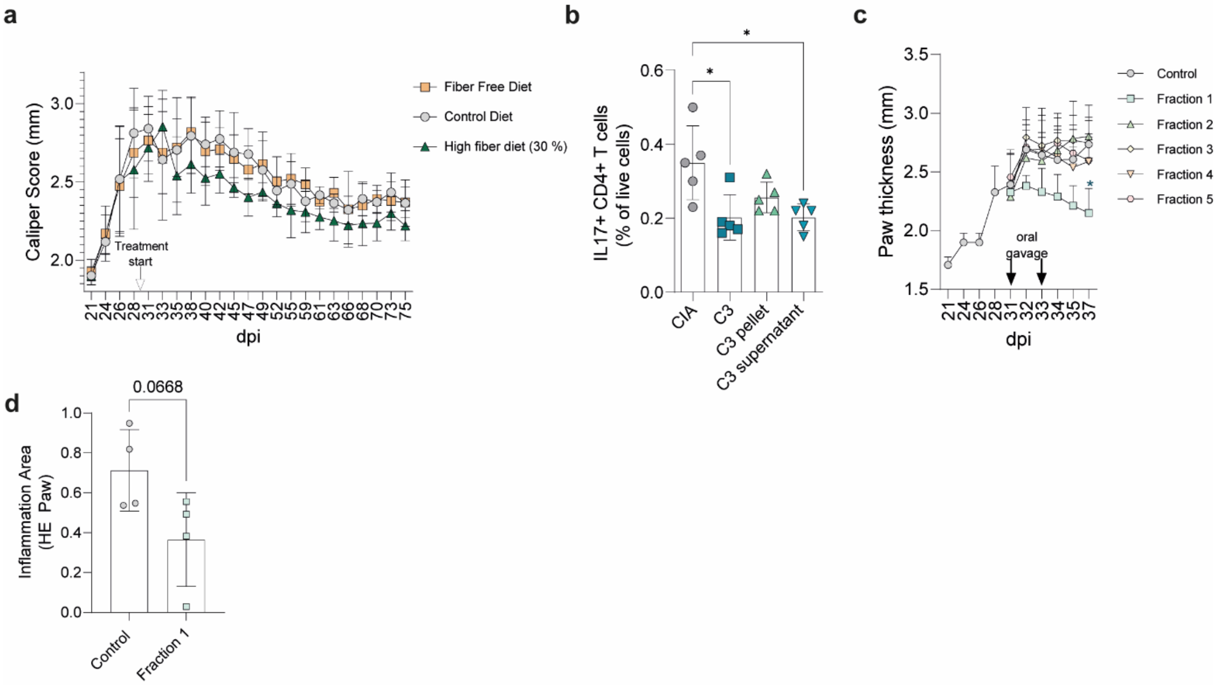


**Supplementary Figure 1**

**a.** Clinical arthritis score shown as paw thickness (mm) of CIA mice on control diet, a high fiber diet (30 % fiber) or a fiber free diet starting 30 dpi **b.** IL-17+ CD4+ T cells in the spleen of CIA mice treated with FMT of naïve donors, C3-treated donors, supernatant of C3-treated donors or pellet of C3-treated donors **c.** Clinical arthritis score shown as paw thickness (mm) of CIA mice receiving different parts of the FMT supernatant separated by size exclusion chromatography by oral gavage **d.** Inflammation area in the paws of CIA mice ± Fraction 1. Data are expressed as the mean ± sd. Statistical difference was determined by One-way ANOVA (b) and t-test (d). *$p < 0.05$; dpi = days post immunization

### Sup. Fig. 2

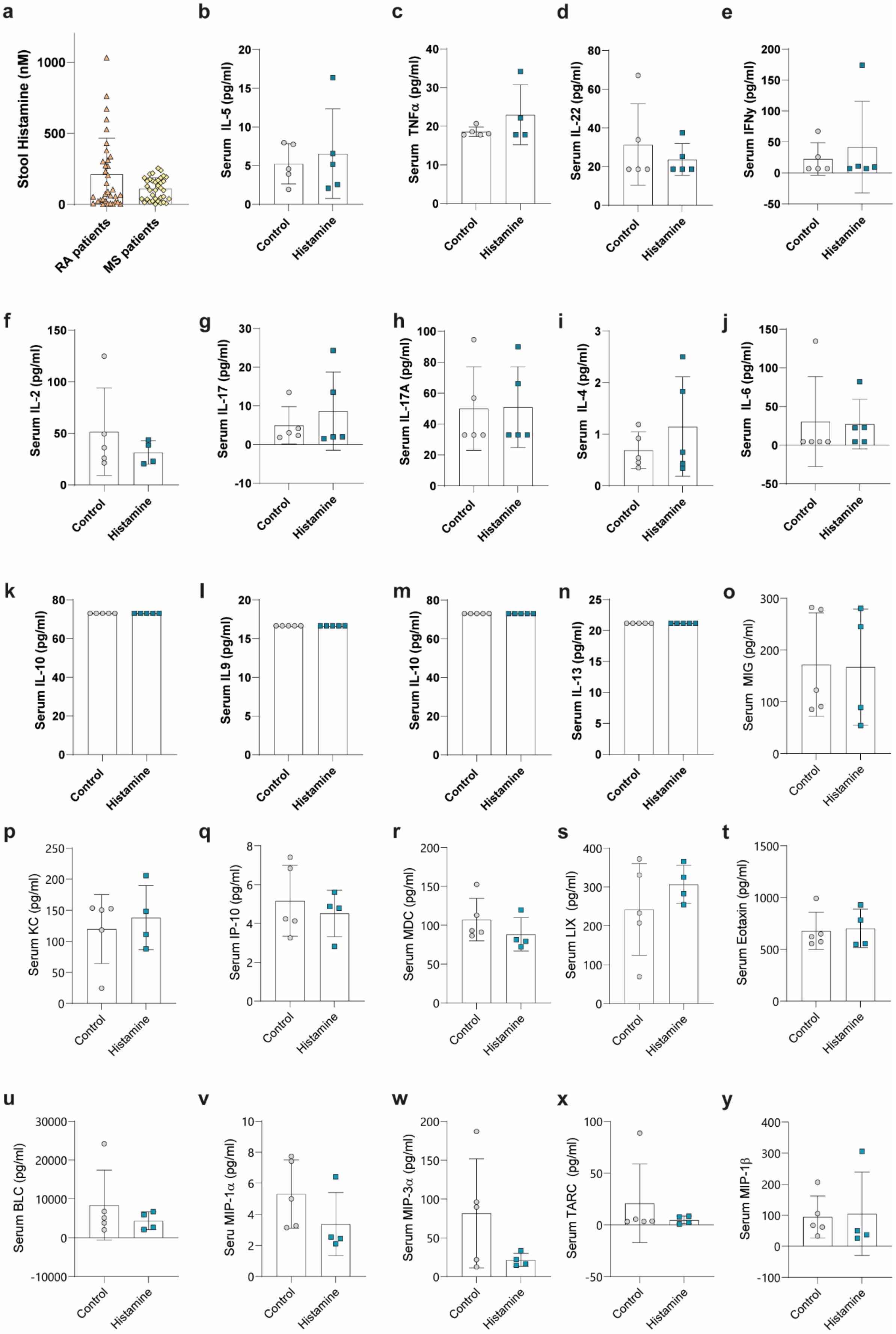

a
Stool Histamine (nM)
RA patients
MS patients
b
Serum IL-5 (pg/ml)
c
Serum TNFα (pg/ml)
d
Serum IL-22 (pg/ml)
e
Serum IFNγ (pg/ml)
f
Serum IL-2 (pg/ml)
g
Serum IL-17 (pg/ml)
h
Serum IL-17A (pg/ml)
i
Serum IL-4 (pg/ml)
j
Serum IL-6 (pg/ml)
k
Serum IL-10 (pg/ml)
l
Serum IL9 (pg/ml)
m
Serum IL-10 (pg/ml)
n
Serum IL-13 (pg/ml)
o
Serum MIG (pg/ml)
p
Serum KC (pg/ml)
q
Serum IP-10 (pg/ml)
r
Serum MDC (pg/ml)
s
Serum LIX (pg/ml)
t
Serum Eotaxin (pg/ml)
u
Serum BLC (pg/ml)
v
Seru MIP-1α (pg/ml)
w
Serum MIP-3α (pg/ml)
x
Serum TARC (pg/ml)
y
Serum MIP-1β
Control
Histamine

**Supplementary Figure 2**

**a.** Stool histamine levels (nM) of RA and MS patients **b.** Serum IL-5 **c.** Serum TNFα **d.** Serum IL-22 **e.** Serum IFNγ **f.** Serum IL-2 **g.** Serum IL-17 **h.** Serum IL-17A **i.** Serum IL-4 **j.** Serum IL-6 **k.** Serum IL-10 **l.** Serum IL-9 **m.** Serum IL-10 **n.** Serum IL-13 **o.** Serum MIG **p.** Serum KC **q.** Serum IP-10 **r.** Serum MDC **s.** Serum LIX **t.** Serum Eotaxin **u.** Serum BLC **v.** Serum MIP-1α **w.** Serum MIP-3 α **x.** Serum TARC **y.** Serum MIP-1β. Data are expressed as the mean ± sd. Statistical difference was determined by Student's t-test. $*p < 0.05$;

SupFig3

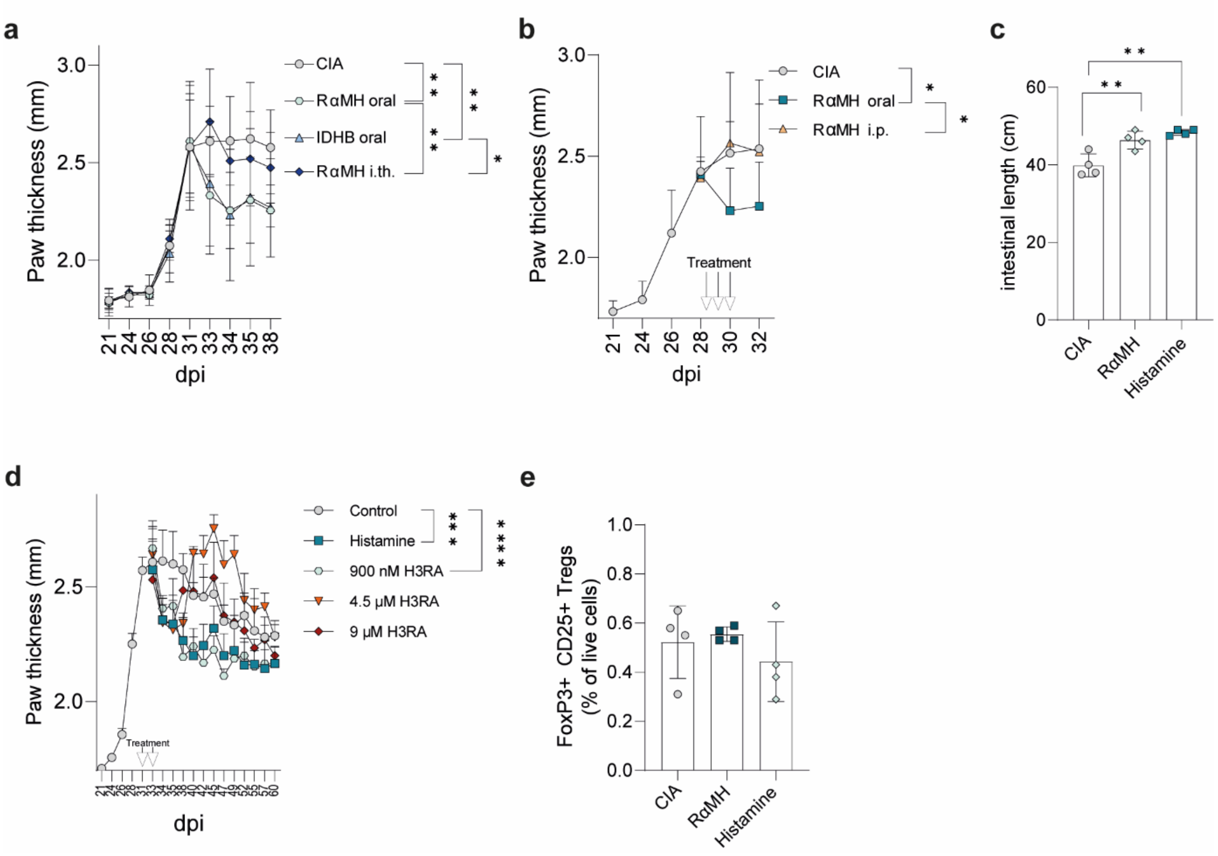


**Supplementary Figure 3**

**a.** Clinical arthritis score shown as paw thickness (mm) of CIA mice treated orally or intrathecally (i.th.) with the H3R agonist RαMH, or orally with the H3R agonist Immethidrine dihydrobromid (IDHB) **b.** Clinical arthritis score shown as paw thickness (mm) of CIA mice treated orally or intraperitoneally with RαMH **c.** intestinal length of CIA mice ± oral treatment with RαMH or histamine **d.** Clinical arthritis score shown as paw thickness (mm) of CIA mice treated orally with histamine or different concentrations of the H3R agonist RαMH. **e.** FoxP3+ CD25+ regulatory T cells in the spleen of CIA mice ± oral treatment with RαMH or histamine. Data are expressed as the mean ± sd. Statistical difference was determined by One way-ANOVA (c, e) of area under the curve (a, b, d). *p < 0.05; **p < 0.01; ***p < 0.001. dpi = days post immunization.

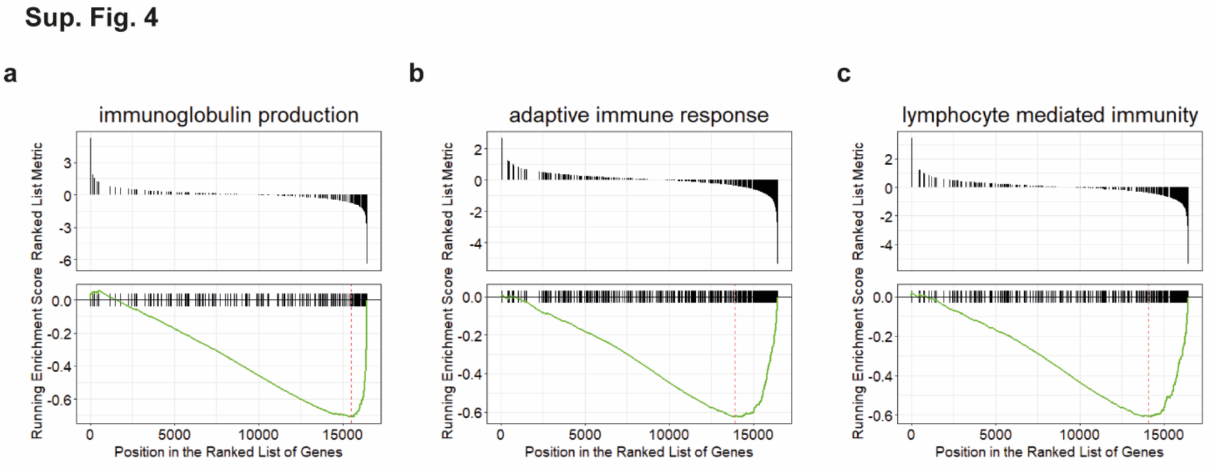


**Supplementary Figure 4**

**a**. Running Enrichment Score and Ranked List Metric of the Immunoglobulin production pathway gene set **b**. Running Enrichment Score and Ranked List Metric of the adaptive immune response pathway gene set **c.** Running Enrichment Score and Ranked List Metric of the lymphocyte mediated immunity pathway gene set.

**Sup. Fig. 5**

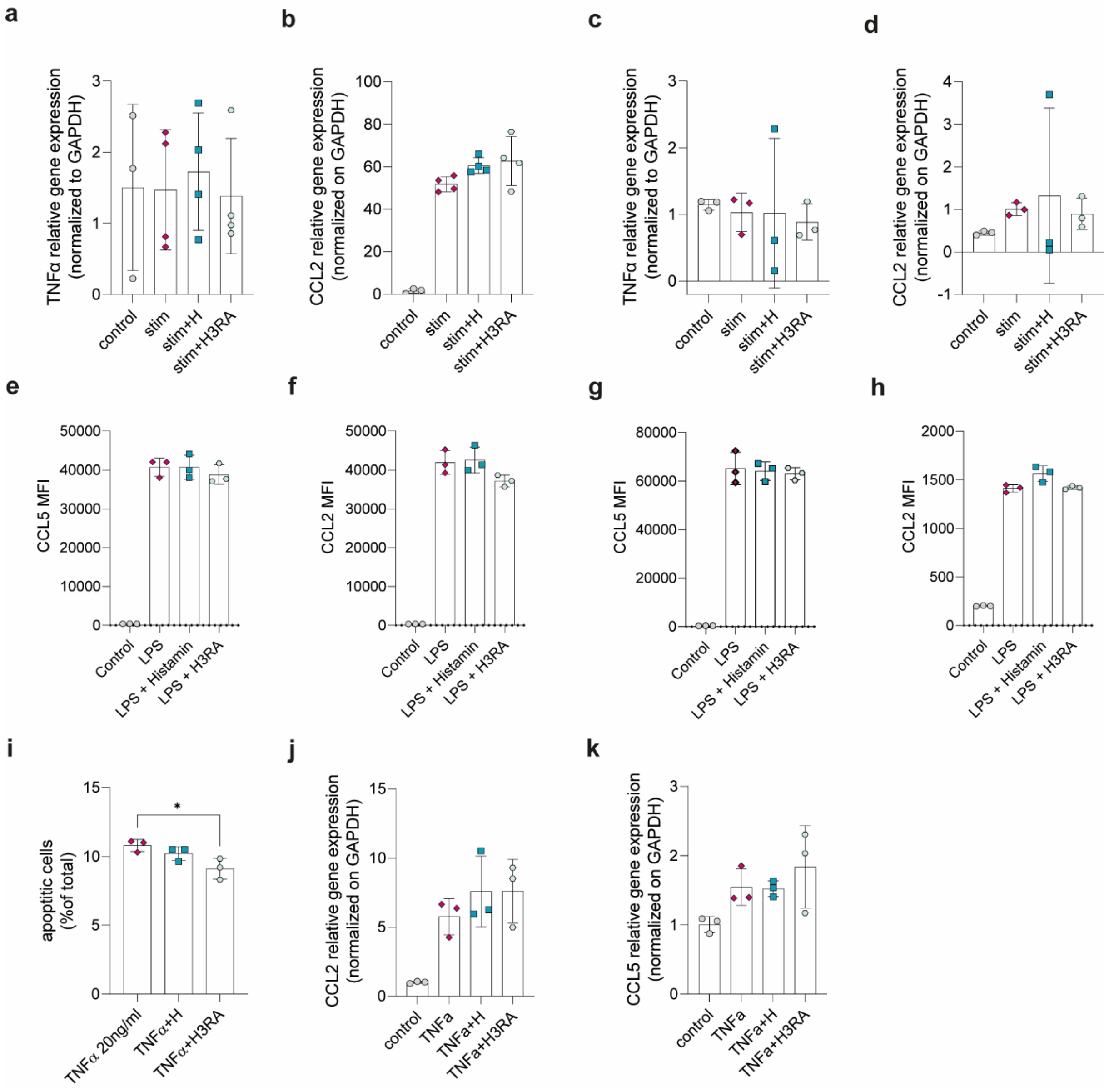


**Supplementary Figure 5**

**a.** Relative TNFα gene expression of astrocytes ± histamine or RαMH **b.** Relative CCL2 gene expression of astrocytes ± histamine or RαMH **c.** Relative TNFα gene expression of microglia ± histamine or RαMH **d.** Relative CCL2 gene expression of microglia ± histamine or RαMH **e.** CCL5 MFI in cell culture supernatant of astrocytes after stimulation with histamine or RαMH **f.** CCL2 MFI in cell culture supernatant of astrocytes after stimulation with histamine or TNFα **g.** CCL5 MFI in cell culture supernatant of microglia after stimulation with histamine or RαMH **h.** CCL2 MFI in cell culture supernatant of microglia after stimulation with histamine or TNFα **i.** apoptotic cells after stimulation of N2A neuronal cell line with TNFα, TNFα + histamine or TNFα +

TNFα **j.** CCL2 gene expression of N2A neuronal cell line with TNFα, TNFα + histamine or TNFα + TNFα **k.** CCL5 gene expression of N2A neuronal cell line with TNFα, TNFα + histamine or TNFα + TNFα. Data are expressed as the mean ± sd. Statistical difference was determined by One way-ANOVA. *$p < 0.05$; **$p < 0.01$; ***$p < 0.001$.

**Sup. Fig. 6**

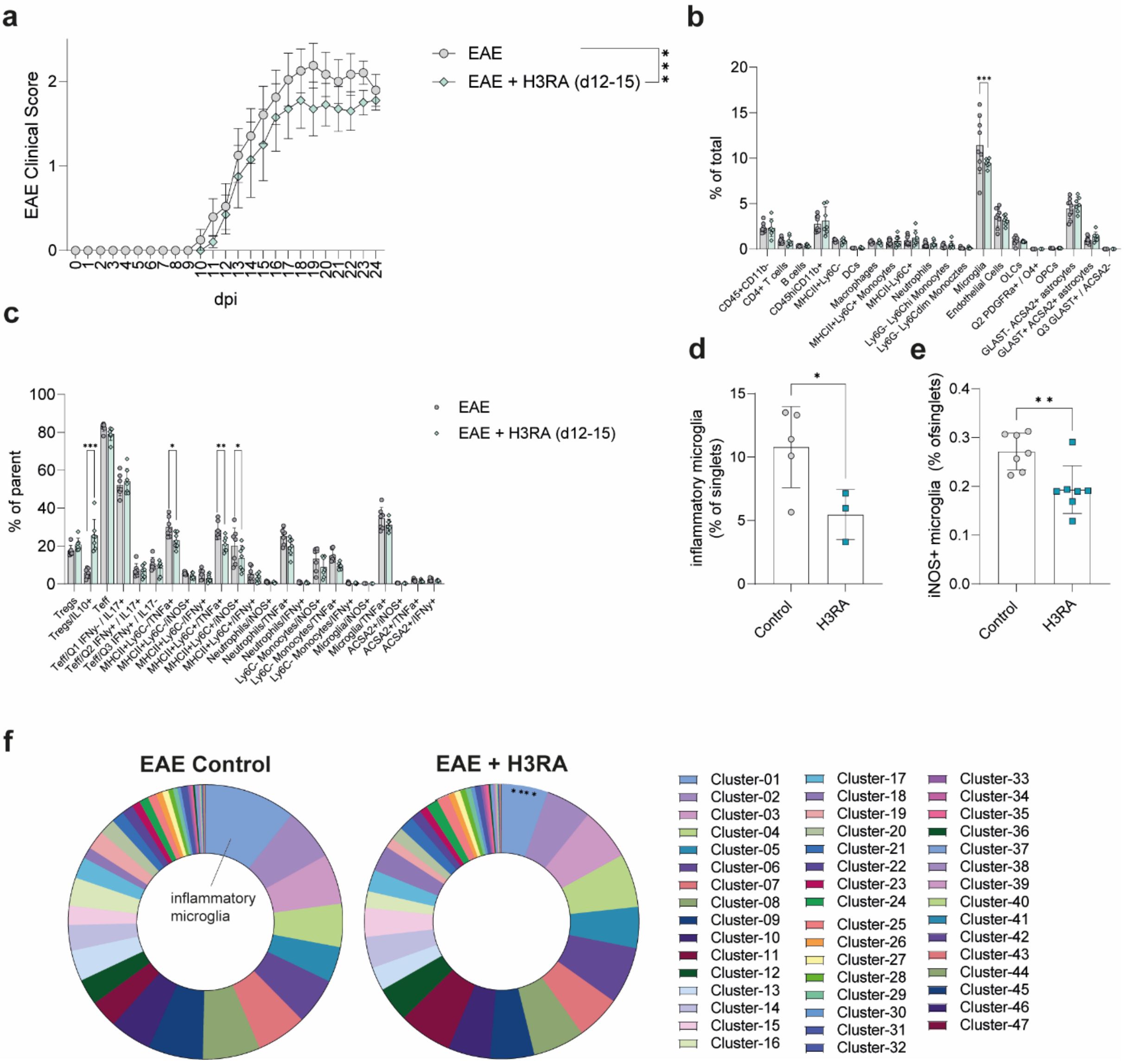


**Supplementary Figure 6**

**a.** EAE clinical score of mice ± RαMH from day 12-14 post immunization **b.** Surface staining of cell populations in the CNS **c.** Intracellular staining of cell populations in the CNS **d.** inflammatory microglia **e.** iNOS+ microglia **f.** Donut representation of the different cell clusters in the CNS after RαMH treatment. Data are expressed as the mean ± sd. Statistical difference was determined by One-way-ANOVA. *p < 0.05; **p < 0.01; ***p < 0.001. dpi = days post immunization

Sup. Fig. 7

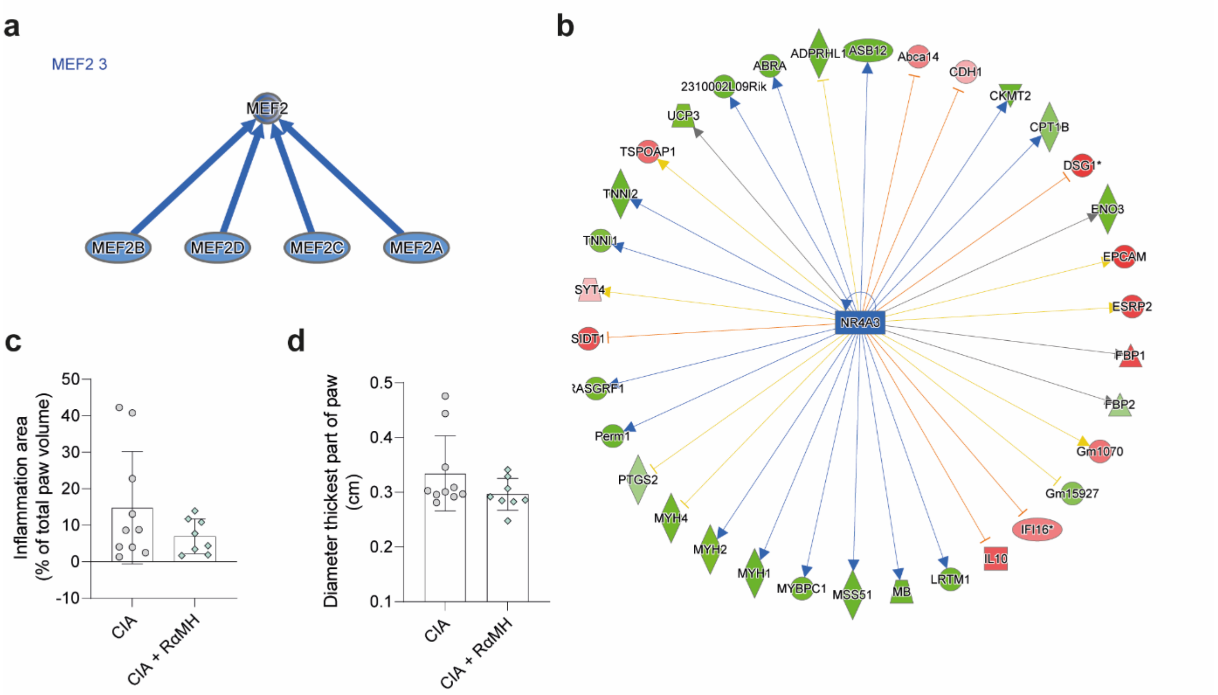


**Supplementary Figure 7**

**a.** Causal Network of NR3A3 Neurovascular Coupling Pathway of RNAseq data from CD11b- cells from nervus plantaris (N.p.) **b.** Causal network of MEF2 Neurovascular Coupling Pathway RNAseq data from CD11b- cells from nervus plantaris (N.p.) **c.** Edema area calculated from MRI T2 STIR of hind paws (% of total paw volume) of CIA mice ± RαMH **d.** Diameter of the thickest part of the paws (cm) calculated from MRI of CIA mice ± RαMH. Data are expressed as the mean ± sd. Statistical difference was determined by Student's t-test.
